# N-terminally acetylated Met11-Tau: a new pathological truncated Tau species with functional relevance in Alzheimer disease

**DOI:** 10.1101/2025.10.03.680200

**Authors:** Sarah Guedjdal, Coline Leghay, Maxime Derisbourg, Sabiha Eddarkaoui, Simon Lecerf, Florian Vermon, Raphaelle Caillierez, Séverine Begard, Claire Regost, Charlotte Laloux, Paulo J. da Costa, Kevin Carvalho, Giovanni Chiappetta, Yann Verdier, Valérie Buée-Scherrer, Vincent Deramecourt, Susanna Schraen, David Blum, Franck Martin, Luc Buée, Malika Hamdane

## Abstract

Neurodegenerative diseases like Alzheimer’s disease (AD) are characterized by progressive accumulation of pathological Tau proteins. Among the diverse Tau species, truncated variants are emerging as key contributors, yet their identity remains elusive, particularly for the N-terminal truncated ones. The present study identifies and characterizes a novel N-terminally truncated and N-alpha-acetylated form of the Tau protein. Using a newly developed antibody specifically targeting this truncated variant, we demonstrate that this species accumulates early in degenerating neurons in both transgenic mouse models of AD-related Tau pathology and post-mortem brain tissues from AD patients. Importantly, *in vivo* functional experiments reveal that expression of this truncated Tau species exacerbates Tau pathology, whereas targeted immunotherapeutic with the specific antibody significantly reduces pathological Tau accumulation and prevents associated memory impairments. These findings position this newly identified Tau variant as both a marker of neurofibrillary degeneration and a pathogenic driver of neurodegeneration and supports its potential as a therapeutic target in Tau-related disorders, notably AD.

## INTRODUCTION

Tau is a microtubule-associated protein, primarily expressed in neurons, where it plays, among others, a fundamental role in microtubules stabilization and intracellular transport. In the human adult brain, six Tau isoforms are produced from a single gene by alternative splicing of exons 2, 3 and 10 (*1*). Tauopathies are a group of neurodegenerative disorders-including Alzheimer’s disease (AD), progressive supranuclear palsy (PSP), and frontotemporal lobar degeneration (FTLD)-characterized by pathological modifications of the Tau protein such as hyperphosphorylation, truncation and aggregation (*2*). AD is the most common tauopathy and the leading cause of dementia. A key neuropathological hallmark of AD is the neurofibrillary degeneration (NFD), which consists in intracellular aggregates of Tau proteins bearing abnormal post-translational modifications. Neuropathological and biochemical studies have consistently shown that the progression of NFD in cortical brain areas strongly correlates with cognitive decline in AD, highlighting Tau as a central player in AD pathology (*3–5*). However, the precise mechanisms underlying the formation and the regional spread of NFD remain incompletely understood. Key unresolved questions include the identification of the particular forms of Tau that confer toxicity and the mechanisms by which these species promote neurodegeneration. Among the diverse Tau species observed in AD brains, truncated variants are increasingly recognized as potential pathogenic contributors (*6–10*). Using an optimized proteomic approach, we previously identified several novel N-terminally truncated Tau species from the AD human brain tissue, including a variant starting at the residue methionine11 (Met11-Tau) (*11*). This Tau variant is particularly interesting because Met11 is located in exon1, the N-terminal region common to all Tau protein isoforms. Although the functional roles of exon1 have not been extensively investigated, emerging evidences suggest that modifications in this region may influence Tau functions and contribute to Tau pathology (*11–14*). The N-terminal domain of Tau is thought to play a structural role, notably through its involvement in the formation of a compact “paperclip” conformation, which results from intramolecular interaction between N-terminal and C-terminal domains (*15–17*). Consequently, the loss of the outermost N-terminus of Tau, as encountered in Met11-Tau, might have significant functional and pathological implications. Notably, further analyses of our original proteomic data (see below) revealed that Met11-Tau is present in an N-terminally acetylated form (AcMet11-Tau). N-terminal acetylation is among the most common protein modifications in eukaryotes, affecting approximately 80% of human proteins (*18, 19*). This irreversible co-translational modification involves the transfer of an acetyl group from acetyl-coenzyme A to the α-amino group of a nascent polypeptide’s N-terminus, catalyzed by Nα-terminal acetyltransferases (NATs) (*20*). The functional implications of N-terminal acetylation are diverse and significant, including the regulation of protein stability and conformation. Moreover, dysregulation of N-terminal acetylation has been involved in a range of human diseases, including cancer and neurodegenerative disorders, underscoring its critical role in maintaining cellular homeostasis (*21, 22*). Given the established functional roles of Tau truncation and N-terminal acetylation, elucidating the contribution of the newly identified AcMet11-Tau species to AD-related Tau pathology is of considerable interest. To address this issue, we generated a monoclonal antibody with high specificity for the AcMet11-Tau variant. Through studies in a transgenic mouse model of tauopathy and analyses of postmortem human brain tissues, we demonstrated the presence of AcMet11-Tau in neurons displaying NFD at early pathological stages. Notably, immunotherapy targeting AcMet11-Tau ameliorated both pathological burden and memory deficits in Tau transgenic mice. Collectively, these findings identify AcMet11-Tau as a previously unrecognized, early pathological Tau species with physiopathological and therapeutic implications for AD.

## RESULTS

### Identification of AcMet11-Tau and generation of a specific antibody

By combining immunoprecipitation (IP) and capillary liquid chromatography-tandem mass spectrometry (LC-MS/MS), we previously identified 18 amino-truncated Tau species in human brain samples (*11*). In the present study, we have reanalyzed these proteomic data with a focus on post-translational modifications and identified a Tau variant starting at Met11 residue, bearing an N-terminal acetylation hereafter referred to as AcMet11-Tau (Fig. 1A). To validate this finding and to investigate the pathophysiological relevance of AcMet11-Tau, we developed a monoclonal antibody specifically targeting this variant, termed 2H2D11 (Fig. 1B). Indirect Elisa, using different Tau peptides, showed 2H2D11 to be specific for AcMet11-Tau peptide (Fig. 1C). Notably, this antibody did not recognize either the unmodified Met11 (Met11-Tau peptide) or the non-truncated Tau peptide containing N-α-acetyl-Methionine in a different Tau sequence context (AcMet1-Tau peptide), thereby confirming its high sequence and modification specificity. To further validate the specificity of 2H2D11 antibody, we generated stable SH-SY5Y neuroblastoma cell lines by using tetracycline inducible expression vectors containing the coding sequence of either Full length (FL-Tau) or Met11-Tau. Western blot analysis revealed that 2H2D11 detected Tau expression exclusively in Met11-Tau-expressing cells (hereafter referred to as AcMet11-Tau cells), with no signal in FL-Tau cells (Fig. 1D). Specificity was further confirmed by peptide blocking experiments, where pre-incubation of 2H2D11 with the AcMet11-Tau peptide abolished the signal. Additionally, a sandwich Elisa using 2H2D11 demonstrated strong immunoreactivity only in extracts from AcMet11-Tau-expressing cells, while FL-Tau cell extracts yielded signals comparable to blank controls (Fig. 1D, bottom). Collectively, these results establish 2H2D11 antibody as a highly specific and reliable tool for the detection of the AcMet11-Tau variant, providing a valuable resource for further investigation of its pathophysiological role.

**Fig. 1.**
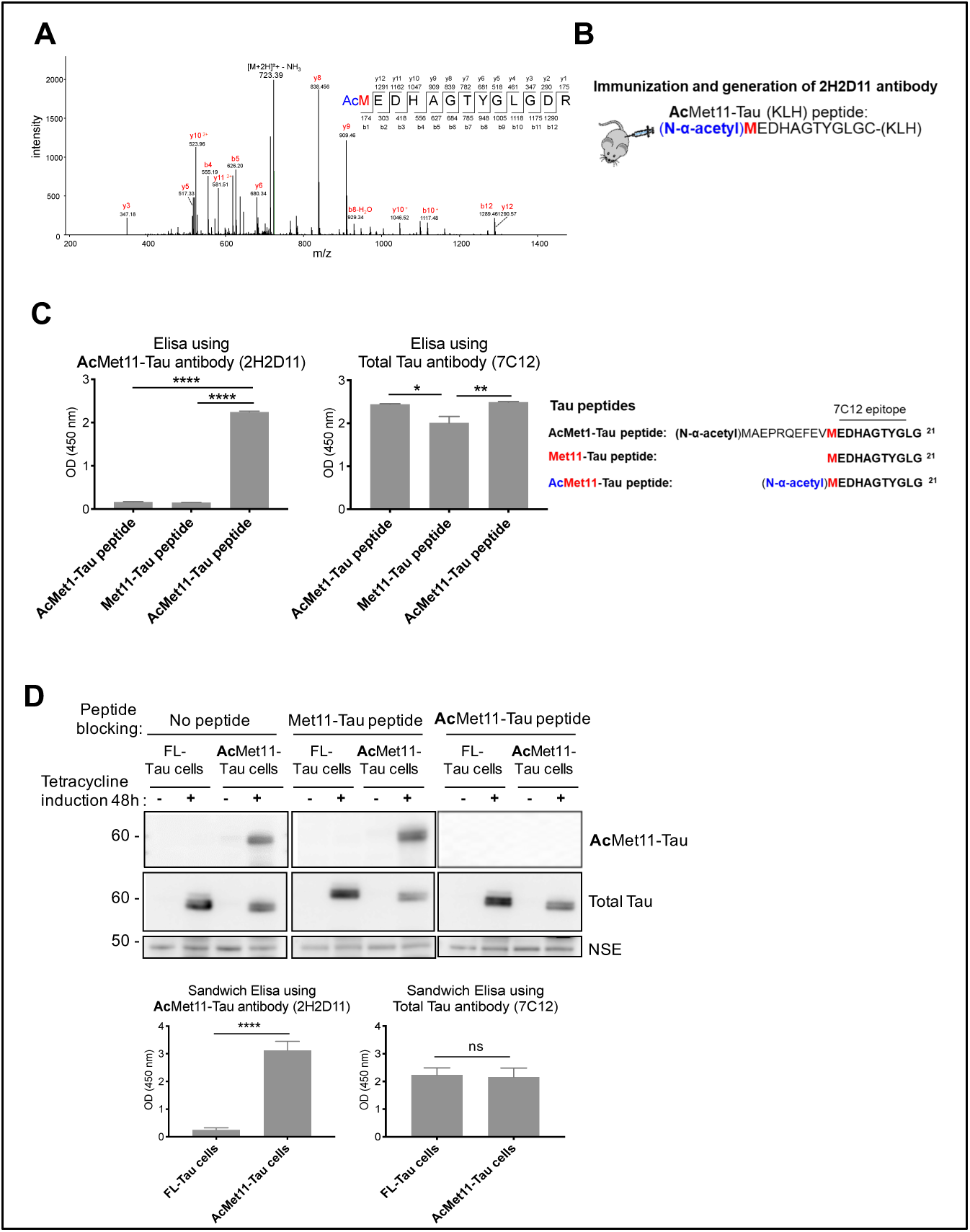
Discovery of AcMet11-Tau protein and validation of its antibody specificity. (**A**) MS/MS spectra of N-terminally acetylated peptide Met11-Tau from a human brain (Braak stage III, table S1). Charge:+2. Monoisotopic m/z: 732.31 Da; MH+: 1463.62.RT: 27.86. Scan 154. G140418_0427_c_ymv.raw Identified with Mascot v1.30; ion score = 54; exp value: 3^E^-003. (**B**) Representation of 2H2D11 antibody production with the peptide sequence used for immunizations. (**C**) Histograms indicated representative Elisa OD values obtained by 2H2D11 and 7C12 (Total-Tau) antibodies, (mean ± SEM, n=3 independent experiments, *p= 0.0125, **p = 0.0077, **** p<0.0001, Ordinary one-way ANOVA followed by Fisher’s LSD post hoc test). Amino Acids sequences of the synthetic peptides used in Elisa are shown, the line indicates the 7C12 epitope, according to the longest Tau isoform. (**D**) Protein extracts from SH-SY5Y inducible cell lines overexpressing either Full-length Tau (FL-Tau) or AcMet11-Tau, after 48h of tetracycline treatment, were analysed by: (top) Western blot using 2H2D11 antibody pre-adsorbed or not with Tau peptides; membranes were re-probed with Total-Tau antibody (Tau-Cter) and NSE as loading control; (bottom) Sandwich Elisa; Histograms indicated OD values, (mean ± SEM, n=4 independent experiments, **** p<0.0001, unpaired t test).

### AcMet11-Tau is an early pathological marker in a mouse model of Tau pathology

To investigate a potential association between AcMet11-Tau and NFD, we analyzed brain sections from 12-month-old Thy-Tau22 mice, a model that progressively develops hippocampal tau pathology from 3 months of age and exhibits memory impairments by 7 months (*23, 24*). Immunohistochemical staining with the 2H2D11 antibody revealed no detectable signal in wild-type littermate mice (Wt), whereas Thy-Tau22 mice showed robust hippocampal labelling. Notably, 2H2D11 specifically localized to neurofibrillary tangles (NFT)-like inclusions and neuritic processes (Fig. 2A).

**Fig. 2.**
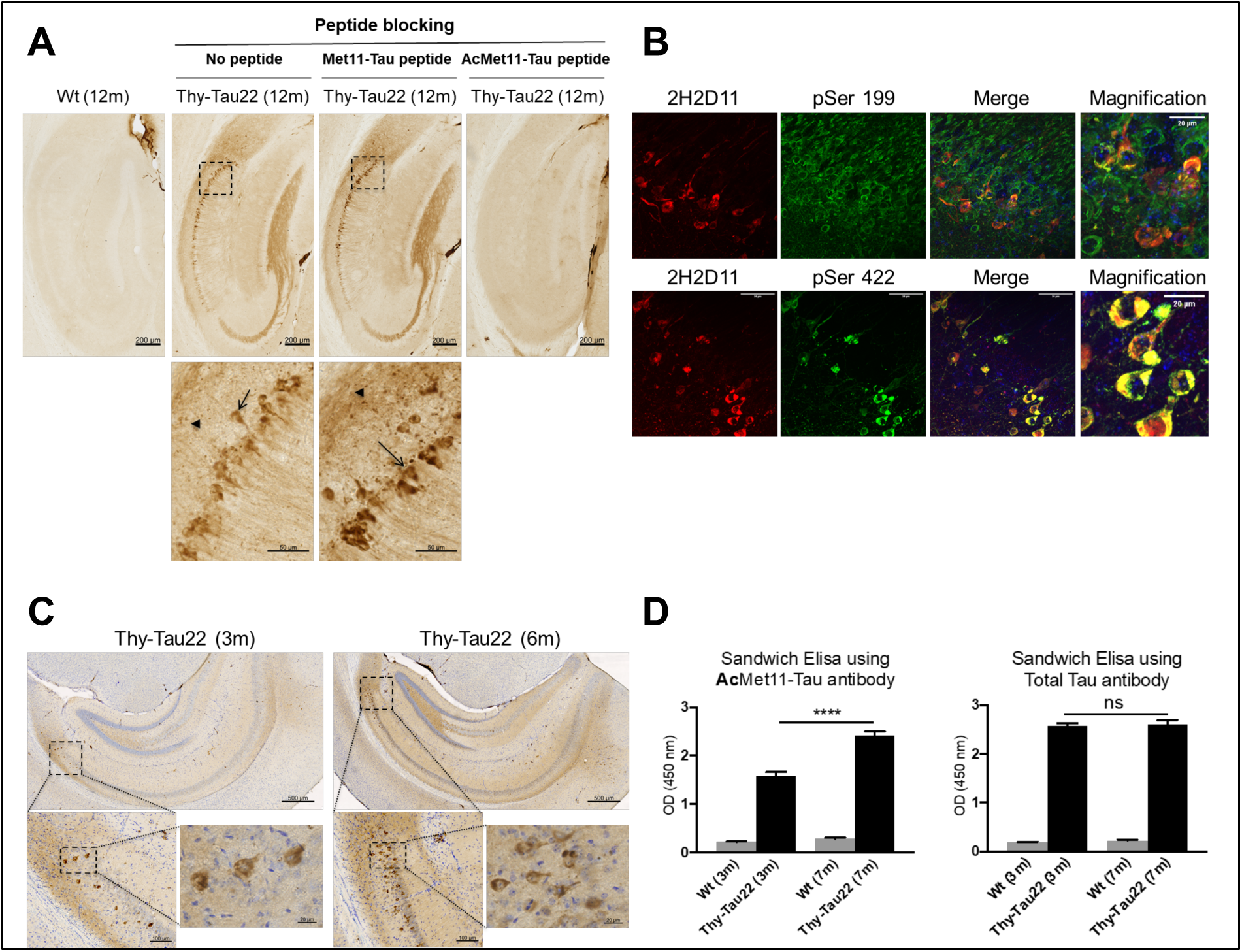
AcMet11-Tau is an early pathological species in Thy-Tau22 mice. (**A**) Representative immunohistochemical analysis of coronal brain sections, using 2H2D11 antibody that labels neurons bearing neurofibrillary tangles (arrow) and neuritic processes (arrowhead) in hippocampus of 12-month-old Thy-Tau22 compared to littermate Wt; blocking experiments with Tau-peptides indicate the specificity of AcMet11-Tau immunostaining. Scale bars = 200 μm and 50 μm (Inset magnification, bottom). (**B**) Representative confocal images of double immunostaining using 2H2D11 antibody with either pSer199 (Top) or pSer422 (bottom) in the hippocampal CA1 region of 9-month-old Thy-Tau22 mice. Scale bars = 50 μm and 20 μm (Merge magnification). (**C**) Immunohistochemical labelling with 2H2D11 of coronal brain sections from 3 and 6-month-old Thy-Tau22 mice. Scale bars = 500 μm (top), 100 μm and 20 μm (bottom). (**D**) 2H2D11 antibody-based Elisa using hippocampal proteins of Thy-Tau22 and their littermate Wt mice of 3 and 7 months of age. Histograms indicated OD values means ± SEM (n=5/group). **** p<0.0001, one-way ANOVA followed by a post hoc Fisher’s LSD test.

To confirm signal specificity, we performed peptide-blocking assays. Pre-incubation of 2H2D11 with the AcMet11-Tau peptide abolished immunoreactivity, whereas pre-incubation with the non-α-acetylated Met11-Tau peptide had no effect (Fig. 2A). These findings support the α-acetylation and sequence specific recognition of AcMet11-Tau *in situ*. Furthermore, we performed double labelling experiments using 2H2D11 with antibodies specific to phosphorylated Tau. The pSer199 antibody broadly labels Tau in neurons whereas the pSer422 antibody specifically detects abnormally phosphorylated pathological Tau in neurons with NFT. Co-labeling revealed that the 2H2D11 antibody selectively labels neurons that are positive for pSer422, but not all pSer199-positive neurons (Fig. 2B) indicating that AcMet11-Tau is preferentially associated with neurons undergoing neurofibrillary degeneration. Strikingly, AcMet11-Tau was detected during early stages of disease progression, largely ahead to the onset of memory deficits. Indeed, immunohistochemical analyses of hippocampal brain sections (Fig. 2C) as well as sandwich Elisa using hippocampal protein extracts (Fig. 2D) demonstrated the presence of AcMet11-Tau as early as 3 months of age in Thy-Tau22 mice. These findings suggest that AcMet11-Tau emerges early in the pathological cascade and may contribute or serve as an early biomarker of Tau pathology.

### AcMet11-Tau is a pathological species specifically detected in the AD brain areas along pathology progression

To assess the presence of AcMet11-Tau in human brain tissue from patients with AD, we first analyzed protein extracts from the temporal cortex and hippocampus of control subjects and AD cases (Braak stages IV-VI), using a 2H2D11-based sandwich Elisa assay. AcMet11-Tau was significantly detected in AD samples as compared to matched controls (Fig. 3, A and C) while total Tau is similarly detected between the two groups (Fig. 3, B and D). To examine disease specificity, across tauopathies, we further analyzed frontal cortex of patients with AD along with patients presenting with PiD and PSP, two primary tauopathies. 2H2D11 immunoreactivity clearly discriminates AD cases from the two other tauopathies (Fig. 3, E and F). Since Tau pathology in PSP patients is more predominant in the mesencephalon than in frontal cortex (fig. S1), we further analyzed AcMet11-Tau expression across two PSP brain regions. AcMet11-Tau remained undetectable, even in the region with robust Tau pathology (Fig. 3, G and H), indicating that AcMet11-Tau is not a feature of PSP. Moreover, we investigated whether AcMet11-Tau is an early pathological marker in AD. To do this, we conducted Elisa-based assay in samples from different brain Brodmann areas (BA) of individuals spanning Braak stages 0 to VI. Prior to analysis, all brain samples were biochemically characterized by immunoblotting to confirm the typical Tau pathology profile, allowing the validation of Braak stages (fig. S2). In BA-38 (anterior temporal pole, affected from Braak stage III), while no significant differences were observed in the total Tau levels detected by 7C12 antibody which recognizes total Tau across the different groups (Fig. 4A), analysis revealed a significant detection of AcMet11-Tau immunoreactivity in Braak stage III-IV and V-VI groups as compared to the Braak stage 0-II group (Fig. 4E). In BA-10 (frontal pole, affected from Braak stage V) and BA-8/9 (dorsolateral prefrontal cortex, affected in later stages), AcMet11-Tau was significantly detected only in the Braak stage V-VI group (Fig. 4F). As expected, AcMet11-Tau detection in BA-17 (primary visual cortex, relatively spared even at the late stages of AD) was not significant (Fig. 4H). A similar detection pattern, consistent with Braak stages and brain regions, was observed using a reference Elisa analysis targeting pathological hyperphosphorylated Tau at pSer396 (Fig. 4, I to L). Importantly, a significant positive correlation was found between optical density values obtained with AcMet11-Tau antibody and those obtained with pSer396 antibody (Fig. 4M). Collectively, these findings indicate that the AcMet11-Tau protein represents an AD disease specific Tau species, closely linked to the pathological Braak staging. Its enrichment in vulnerable cortical regions in AD underscores its potential diagnostic and biomarker utility in AD.

**Fig. 3.**
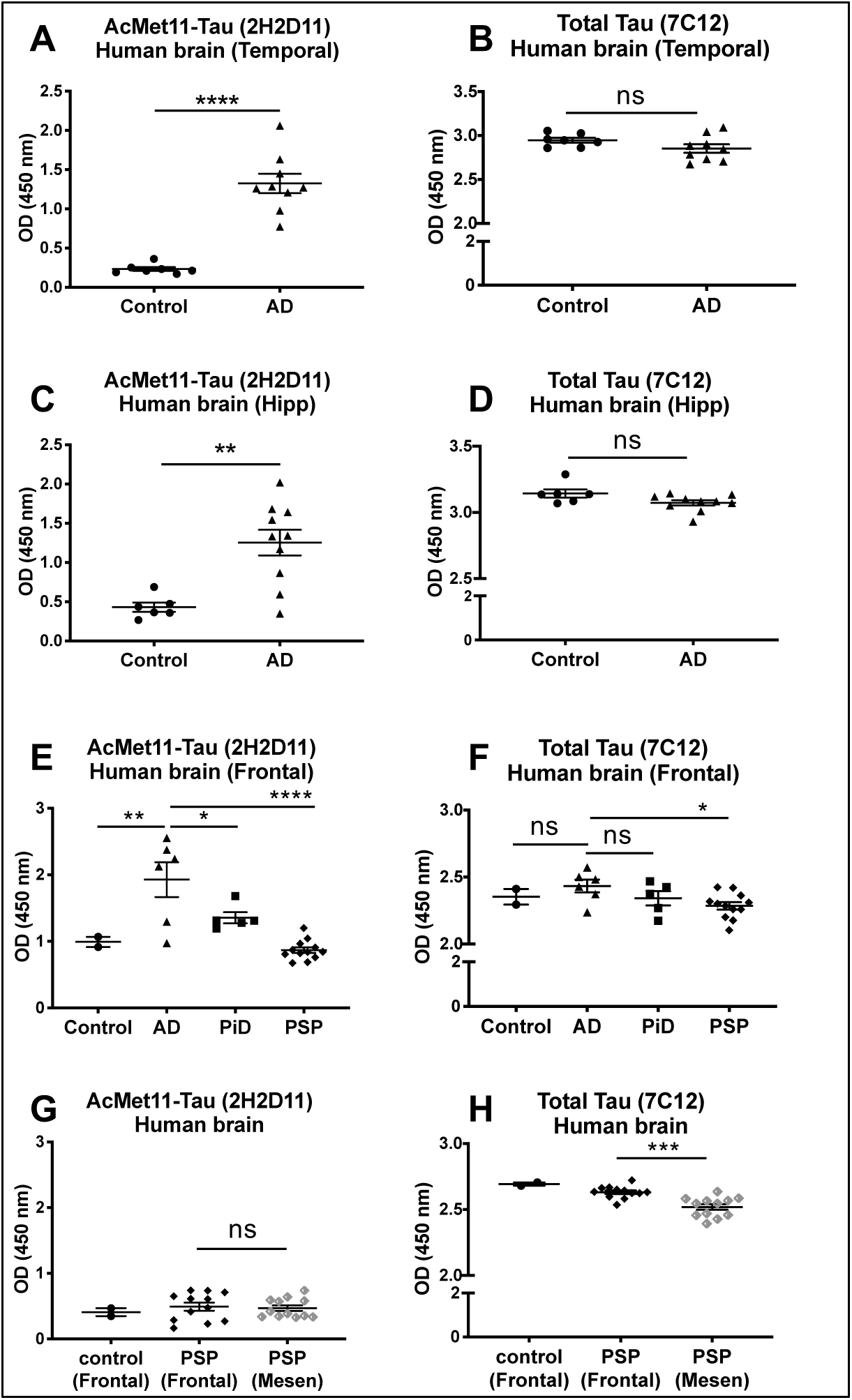
AcMet11-Tau is distinctively detected in AD brains. (**A** to **D**) Protein extracts from temporal and hippocampal regions of controls (n=7 and 6) and AD cases (n=9 and10) were used in 2H2D11 and 7C12 antibodies-based sandwich Elisa (OD values mean ± SEM, **p= 0.0022, ****p<0.0001, unpaired t test). (**E** and **F**) Protein extracts from frontal region of controls (n=2), AD (n=6), PiD (n=5) and PSP cases (n=12) were used in 2H2D11 and 7C12 antibodies-based sandwich Elisa (OD values mean ± SD, **p=0.0029 (AD vs. control), *p=0.0113 (AD vs. PiD), ****p<0.0001(2H2D11, AD vs. PSP), *p=0.0104 (7C12, AD vs. PSP), Ordinary one-way ANOVA followed by Fisher’s LSD post hoc test). (**G** and **H**) Protein extracts from mesencephalon and frontal regions of PSP cases (n=12) were used in 2H2D11 and 7C12 antibodies-based sandwich Elisa (OD values mean ± SEM, ***p=0.0001, Ordinary one-way ANOVA followed by Fisher’s LSD post hoc test).

**Fig. 4.**
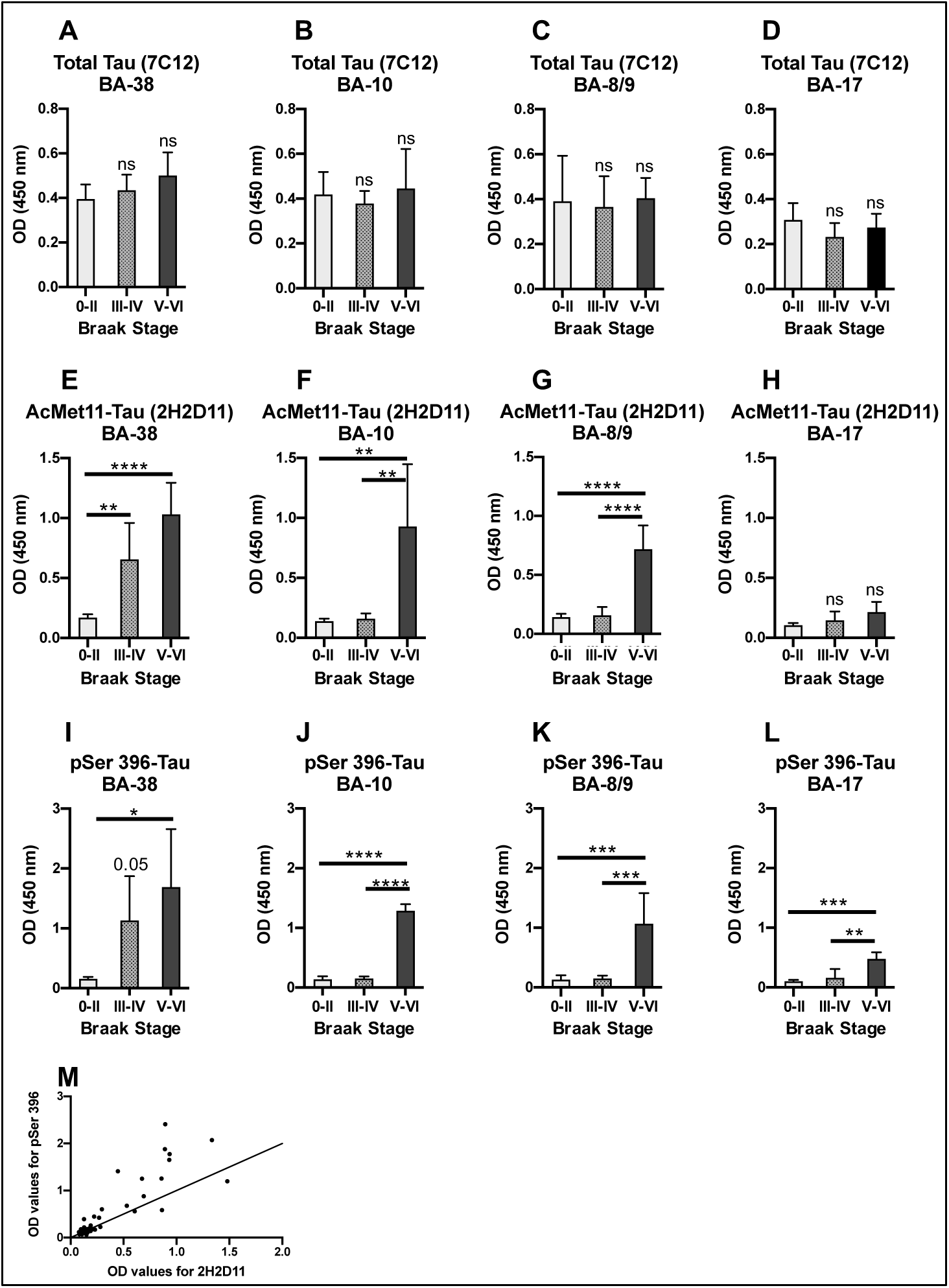
AcMet11-Tau exhibits regional association with Tau pathology throughout AD progression. (**A** to **L**) Elisa Analysis of tissues from different brain areas and individuals at various stages of NFD progression. (**A** to **D**) 7C12 antibody (total Tau) is used as a control for Tau protein detection within the samples. (**E** to **H**) AcMet11-Tau form is detected by 2H2D11 antibody. (**I** to **L**) Phosphorylated Tau is detected using the pSer396 antibody. Data are expressed as the mean optical density ± SD (n=6 (stages 0-II), n=5 (stages III-IV), n=3 (stages V-VI)). *p<0.05, **p<0.01, ***p<0.001, ****p<0.0001, using a one-way ANOVA analysis followed by a Tukey HSD post-hoc test (**M**) Correlation analysis of Elisa OD values for 2H2D11 antibody (x-axis) and pSer 396 antibody (y-axis). Each point corresponds to a sample (one per brain area studied, i.e. four per patient). Significant positive correlation (ρ=0.77 [0.64-0.86], p<0.0001) between OD values obtained with 2H2D11 and pSer396, determined by Spearman’s coefficient test.

### AcMet11-Tau potentiates Tau pathology development in a transgenic mouse model of Tau pathology

To evaluate the AcMet11-Tau pathogenicity, we compared the impact of its overexpression vs. full-length Tau protein (FL-Tau) in the brain of Tau transgenic mice using lentiviral vectors (Lv) designed for neuron-specific expression (*25*). We have performed stereotaxic injections of Lv into the hippocampus of one-month-old Tau transgenic mice, a developmental stage at which Tau pathology is absent (*26*). Prior to stereotaxic injections, we confirmed that both Lv batches drive comparable level of expression in primary neuronal cells, and that Met11-Tau was expressed as N-α-acetylated form (fig. S3). Two months post injections, immunohistochemical analysis of brain sections using an anti-total Tau antibody confirmed similar expression across both experimental groups (Fig. 5, A and B). As anticipated, AcMet11-Tau immunoreactivity was weak in transgenic animals injected with PBS, taken as control, or Lv driving the expression of FL-Tau (Lv-Tau) but was robustly detected in the hippocampi of mice receiving Lv-Met11 (Fig. 5, C to E). Notably, the accumulation of aggregated and fibrillary Tau forms, assessed using AT100 immunostaining, was modestly elevated in mice injected with the Lv-Tau, compared to PBS-treated controls. In stark contrast, mice overexpressing the AcMet11-Tau variant (Lv-Met11) exhibited a pronounced accumulation of fibrillary Tau aggregates (Fig. 5, F to H). These findings strongly suggest that the AcMet11-Tau variant is not merely a passive marker of Tau pathology but functions as an active driver, significantly accelerating NFD *in vivo*.

**Fig. 5.**
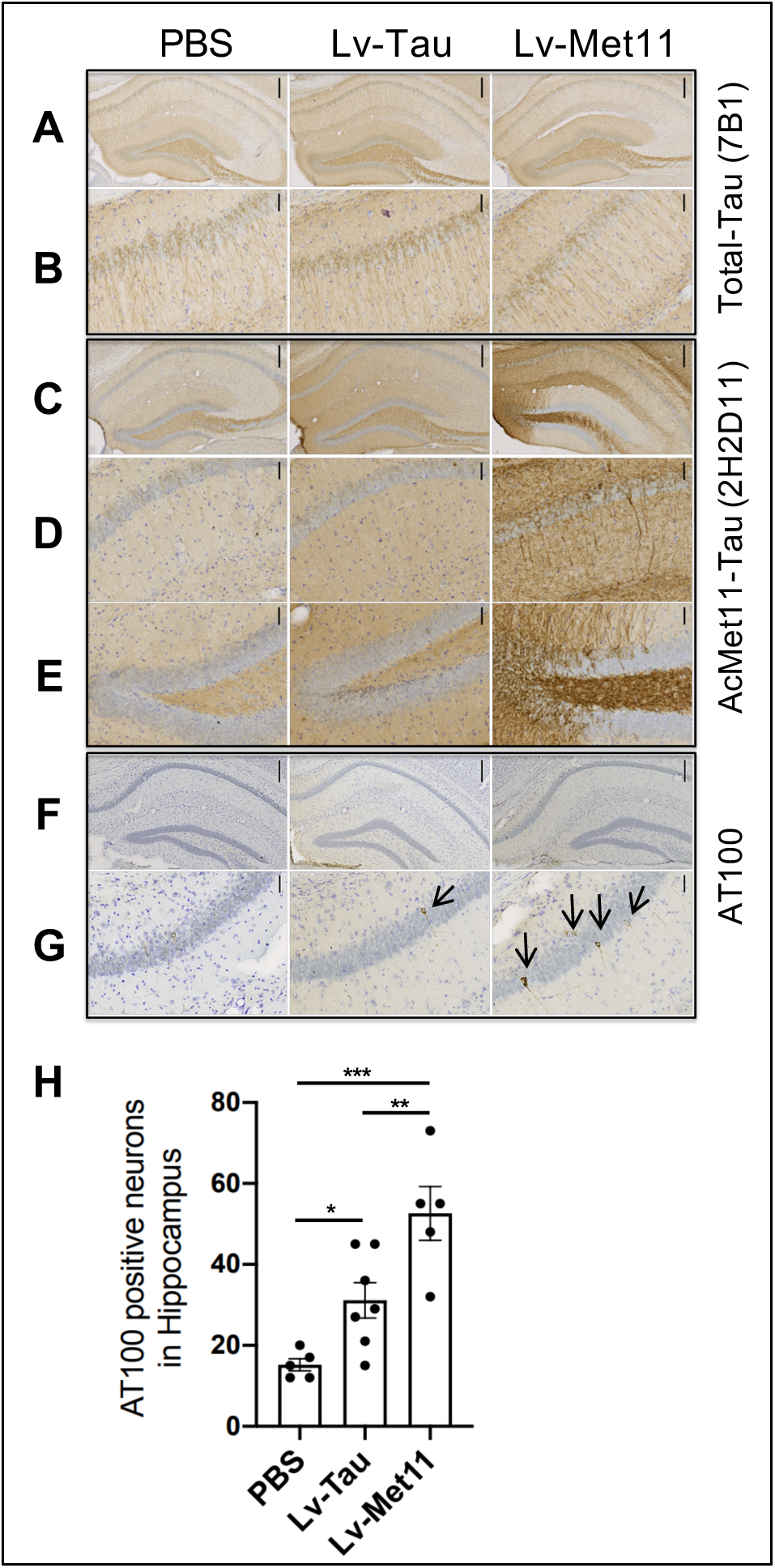
Early expression of AcMet11-Tau in transgenic mice induced an increase in the number of neurofibrillary tangles-bearing neurons. Thy-Tau30 transgenic mice were bilaterally injected with lentival vectors (Lv), into CA1 region of the hippocampus (bregma −2.54. and analyzed 2 months later. (**A** to **G**) Immunohistochemical analysis of coronal brain sections using: (**A** and **B**) Total-Tau antibody (7B1); (**C** to **E**) 2H2D11 antibody; and (**F** and **G**) AT100 that labels neurons bearing NFT (arrows). Panels B, D and G are higher magnification images of CA1 region; panel E is higher magnification images of dentate gyrus. Scale bars = 200 μm (A, C and F), and 50 μm (B, D, E and G). (**H**) Graph represents the number of AT100 positive neurons quantified in 3 sections from left or right hemisphere (bregmas:-1.7 or −1.9, −2.46 or −2,54 and −3.16 or −3.4). Data are expressed as the mean ± SEM of AT100 positive neurons (5-7 hemispheres/group). *p=0.0274, **p=0.0005, ***p=0.0001, ordinary one-way ANOVA followed by Fisher’s LSD post hoc test).

### Beneficial effect of passive immunotherapy against AcMet11-Tau

To investigate whether targeting AcMet11-Tau could confer protective effect against Tau pathology and associated memory deficits, we evaluated a passive immunotherapy approach using the 2H2D11 antibody. Like to a previously described approach (*27*), we have first evaluated the effect of 2H2D11 injection into the hippocampus of Tau transgenic mice. Five days following injection, brain sections were analyzed by IHC using AT100, an antibody that recognizes Tau in its aggregated pathological form, and MC1, a conformational antibody specific to Tau species that precede Tau aggregation and neurofibrillary degeneration. Mice treated with 2H2D11 exhibited a significant reduction of MC1 immunoreactivity as compared to animals injected with a control IgG (Fig. 6, A to C). AT100-positive neurons also appeared to be reduced, albeit not reaching statistical significance within the short treatment window used (Fig. 6, D to F). Given that MC1 detects an earlier stage of Tau aggregation process than AT100 (*28, 29*), our results suggest that immunoneutralization of AcMet11-Tau preferentially impacts early pathogenic Tau species, possibly interfering with the oligomerization process before fibrillary inclusions form. The selective reduction of MC1 staining within just five days highlights the presumable potential of AcMet11-Tau-directed immunotherapy to target upstream, intermediates in the Tau aggregation cascade. To further assess the therapeutic potential of AcMet11-Tau targeting, we then employed a repeated intraperitoneal (IP) immunization approach using the 2H2D11 antibody. Thy-Tau22 mice were injected IP every 15 days, from 3 months i.e. at pathological onset to 7-months of age when Tau pathology is ongoing and memory impairments are present but not maximal in this model. The repeated IP injections were performed for the 2H2D11 antibody or the control IgG (fig. S4). As additional controls, we used littermate Wt mice IP injected with control IgG or PBS, to provide a baseline in behavioral evaluations. Longitudinal body weight monitoring showed no effect of immunization with 2H2D11 as compared to control IgG (fig. S5). Similarly, immunization with 2H2D11 had no impact on animal’s spontaneous motor activity and anxiety-related behavior (fig. S6, A to E). To evaluate the effect of 2H2D11 immunotherapy on memory, we assessed spatial short-term memory using the Y-maze task (Fig. 7A). As expected (*23, 30*), Thy-Tau22 mice injected with control IgG displayed impaired short-term spatial memory, reflected by a lack of preference for the novel arm over the other familiar arm, and a significantly reduced percentage of time spent in the new arm as compared with WT mice. Remarkably, 2H2D11-injected transgenic mice performed similarly to WT mice, showing a significantly increased percentage of time spent in the new arm as compared to Thy-Tau22 mice solely treated with the control IgG, supporting restored spatial memory abilities. To further assess cognitive performance, animals were subsequently assessed in the Barnes maze. The general assessment of spatial learning revealed that all animal groups learned the task by locating the target box over the four days of acquisition (fig. S6F). Although no differences were observed in the path length (fig. S6F), Thy-Tau22 mice injected with control IgG made significantly more primary and total errors as compared to WT littermate controls (Fig. 7, B and C). Notably, Thy-Tau22 mice treated with the 2H2D11 antibody made significantly less errors in reaching the target hole than the Thy-Tau22 mice treated with control IgG, further supporting a beneficial effect of AcMet11-Tau immuno-targeting on spatial learning. In agreement with these behavioral observations, IHC staining of the conformational and pathological Tau AT100 epitope showed a significant decrease of Tau pathology in the hippocampus of Thy-Tau22 mice injected with 2H2D11 antibody (Fig. 8, A and B). We have also assessed some neuroinflammatory markers known to be associated with Tau pathology development in Thy-Tau22 strain (*31, 32*). As expected, our data indicate that these markers increased significantly in the hippocampus of transgenic Tau mice as compared to littermate WT controls (Fig. 8C). Interestingly, Thy-Tau22 mice treated with the 2H2D11 antibody displayed a decrease in the expression of Clec7a, GFAP and Itgax compared to Thy-Tau22 treated with control IgG, supporting the beneficial effect of passive immunotherapy towards neuroinflammatory markers associated to Tau pathology.

**Fig. 6.**
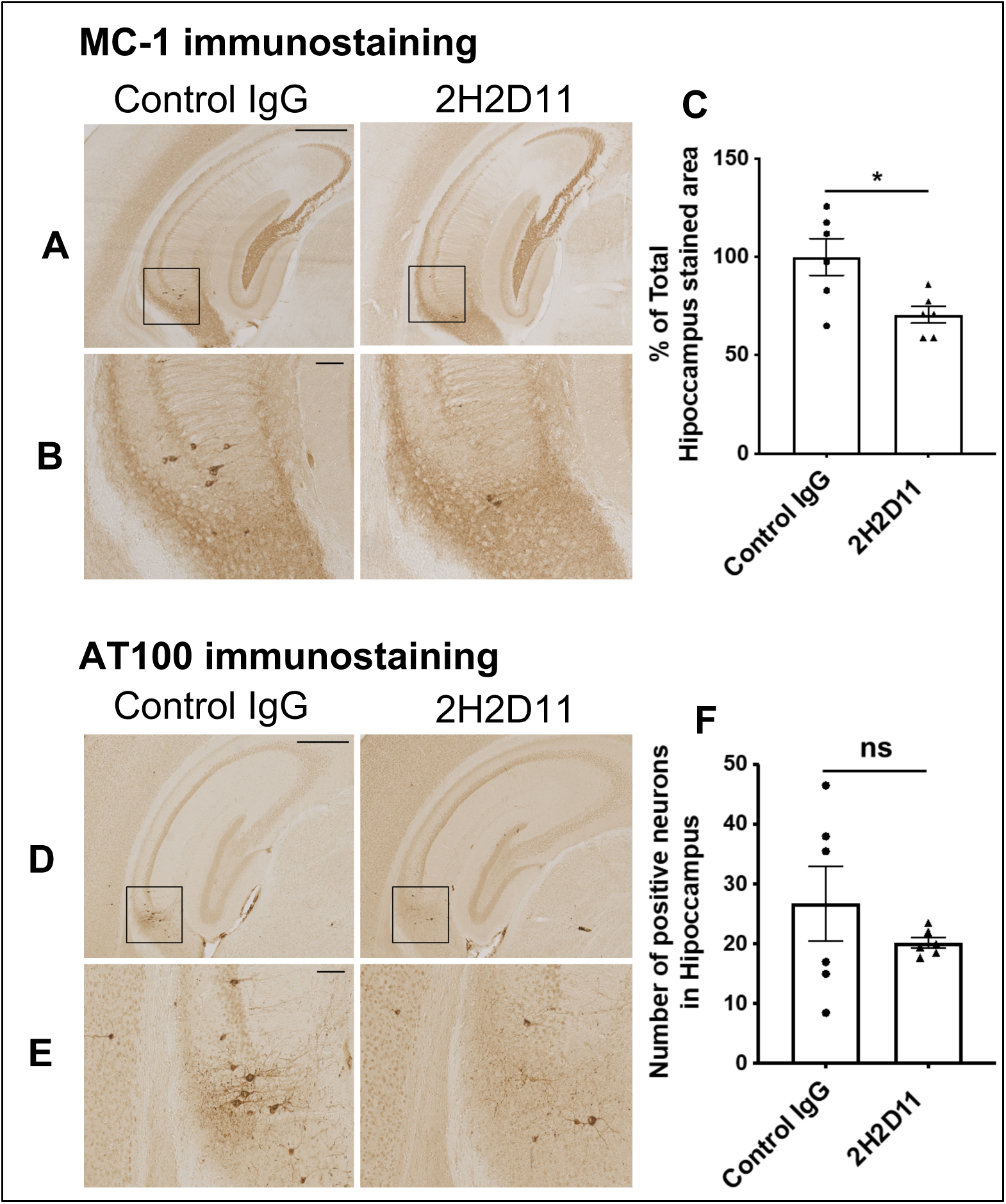
Intracerebral 2H2D11 injection reduces Tau pathology in the hippocampus of transgenic mice. Thy-Tau30 mice (3.5-month-old) received bilateral stereotaxic injections of 2H2D11 or control IgG into the hippocampus. Five days post-injection, brains were processed for immunohistochemistry, using antibodies against pathological Tau protein forms: MC1 and AT100. (**A** and **D**) Representative images of MC1 and AT100 immunostaining, at bregma −2.15 mm, scale bar = 500 μm. (**B** and **E**) Inset magnification, scale bar = 100 μm. (**C** and **F**) MC1-immunostaining, expressed as a percentage of the total hippocampal area and AT100-immunostaing, expressed as the number of positive neurons; quantifications were performed on three coronal sections per animal spanning bregma −1.43 mm to −3.39 mm; Data are presented as mean ± SEM (n = 6 mice per group) and were analyzed using unpaired t-test; 2H2D11 injection significantly reduced MC1 labeled area (*p = 0.0169) and showed a trend toward reduced number of AT100 positive neurons (p = 0.3211).

**Fig. 7.**
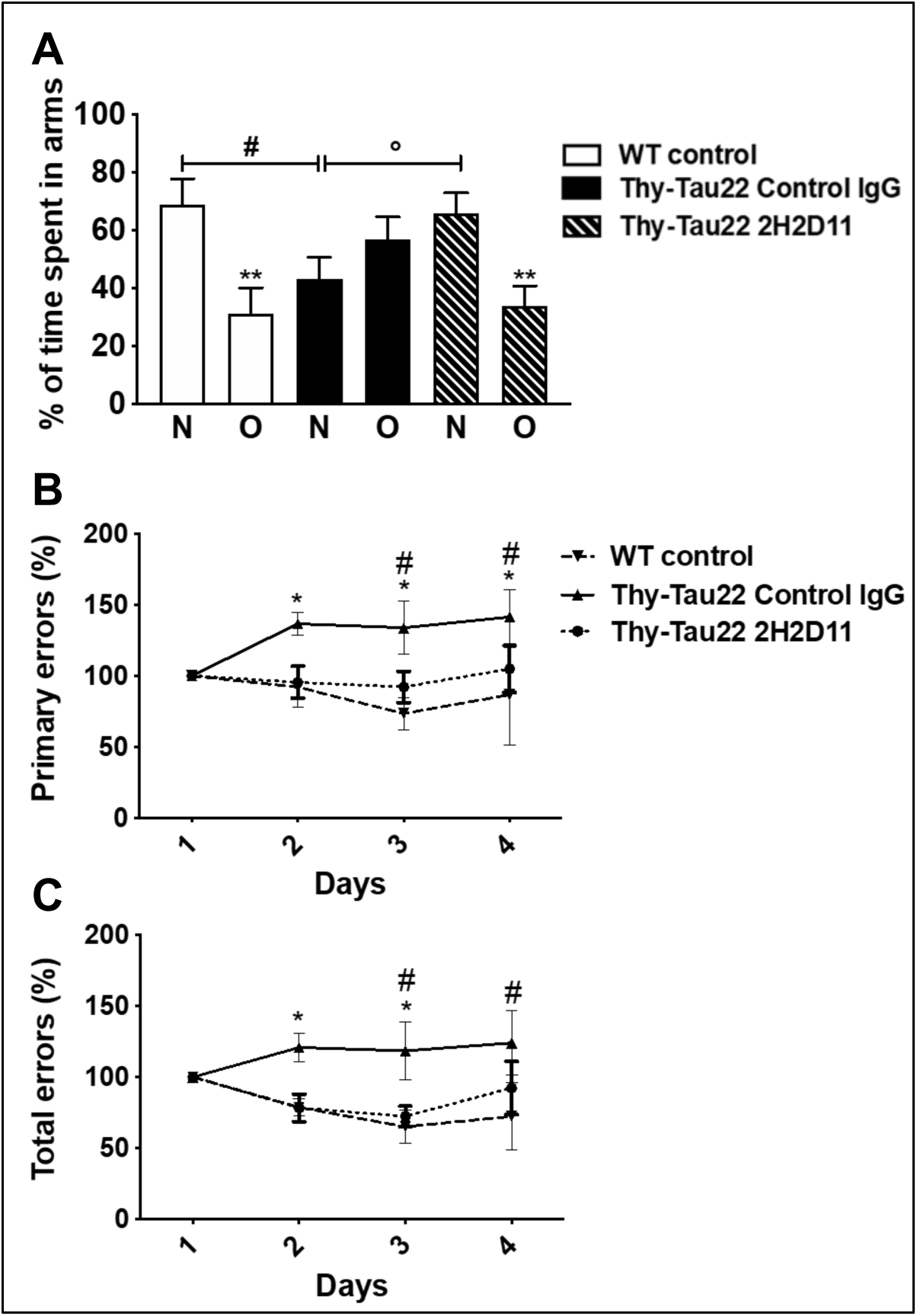
Passive immunotherapy against AcMet11-Tau improves spatial memory and learning of Thy-Tau22 mice. (**A**) In the Y-Maze task, WT control animals exhibited a preference for the novel (N) over the other (O) familiar arm (**p<0.01). Thy-Tau22 treated with control IgG showed an absence of preference for N vs. O (p=0.21) and a percentage of time spent in the N arm significantly reduced as compared to WT (#p<0.05). Conversely, Thy-Tau22 treated with the 2H2D11 antibody exhibited a significant preference for N vs. O (**p<0.01) together with a percentage of time spent in the N arm significantly enhanced as compared to Thy-Tau22 treated with control IgG (°p<0.05). Data were analyzed using One-way ANOVA followed by LSD Fisher test. (**B** and **C**) In the Barnes maze the number of primary and total errors are as expected significantly different between the WT mice and Thy-Tau22 mice treated with control IgG (*p <0.05). Thy-Tau22 mice treated with 2H2D11 antibody made significantly less errors in reaching the target than the Thy-Tau22 mice treated with control IgG (#p <0.05). Data were analyzed using two-away ANOVA followed by a post-hoc HSD Tukey test. All data are presented as mean ± SEM (n= 10-17).

**Fig. 8.**
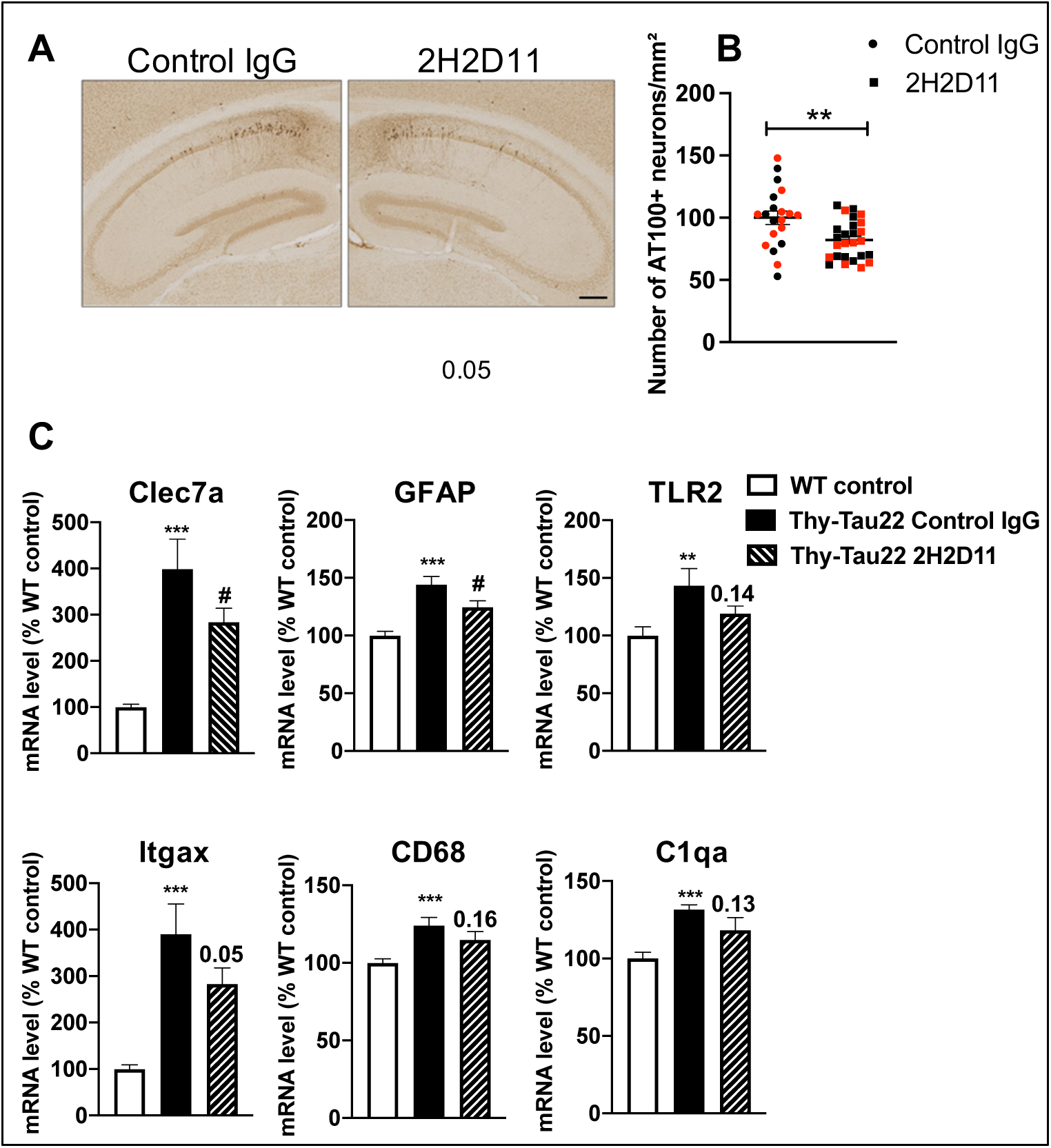
Passive immunotherapy against AcMet11-Tau reduces Tau pathology and its associated neuroinflammatory markers in the hippocampus of Thy-Tau22 mice. (**A**) Representative images of AT100 immunostaining at bregma −2.45 mm, scale bar = 200 μm. (**B**) AT100-immunostaing, expressed as the number of positive neurons per mm^2^, quantifications were performed on three coronal sections per animal spanning bregma −1.43 mm to −3.39 mm; Data are presented as mean ± SEM (n = 20-26 mice per group; females are shown in red) and were analyzed using unpaired t-test; **p < 0.01). (**C**) Treatment with 2H2D11 antibody decreases mRNA expression of Clec7a, GFAP and Itgax (even if its p-value is at the threshold of statistical significance:0.05) in the hippocampus of Thy-Tau22 mice treated with 2H2D11. Thy-Tau22 Control IgG vs. WT, ***p<0.001, **p<0.01; Thy-Tau22 Control IgG vs. Thy-Tau22 2H2D11, #p<0.05; using one-way ANOVA followed by a post hoc Fisher’s LSD test (n = 6/group).

## DISCUSSION

Neurodegenerative diseases termed tauopathies, including Alzheimer’s disease (AD), are defined by the progressive accumulation of pathological Tau species. While post-translational modifications and truncations of Tau have long been associated with its pathological aggregation, the identity of the specific pathogenic driver forms has remained elusive. Here, we identify AcMet11-Tau, an N-terminally truncated and N-α-acetylated Tau variant, as a novel pathological species enriched in early degenerating neurons of both transgenic tauopathy models and AD patient brains. Notably, AcMet11-Tau was absent in the two primary tauopathies analyzed, suggesting a selective association with AD. Using a newly generated specific monoclonal antibody, we demonstrated that AcMet11-Tau is not only a marker of neurofibrillary degeneration, but also an active pathophysiological driver. Overexpression of AcMet11-Tau in transgenic mice exacerbated Tau pathology, while targeted neutralization via passive immunotherapy attenuated both Tau pathology and cognitive deficits. These findings align with prior reports highlighting the role of Tau truncation in tauopathy pathogenesis, particularly in AD (*33*).

While many studies have focused on C-terminally truncated Tau species, N-terminal truncations remain comparatively underexplored (8, 34). Our findings significantly expand current knowledge by identifying an N-terminal truncated Tau variant that is not only novel in sequence but also exhibits an N-α-acetylation, a post-translational modification so far not reported for Tau. To date, the generation of truncated Tau species is traditionally attributed to proteolytic cleavage of FL-Tau (*34*), resulting in fragments which could then contribute to Tau aggregation and neurotoxicity (*7, 8, 33, 35–39*). However, a collection of evidence supports the hypothesis that AcMet11-Tau arises via a non-canonical translation initiation mechanism. First, the AUG codon for Met11 is embedded within an optimal Kozak consensus sequence (*40*), supporting its role as a bona fide alternative translation starting site. Second, Met11 residue is likely acetylated in a co-translational manner by N-acetyl transferase B, an enzyme that is active only when recruited by translating ribosomes (*41*). Such data suggest de novo synthesis rather than post-translational cleavage. Third, our analyses of published ribosome profiling data (*42*) indicate the presence of initiating ribosomes on the Met11 codon, reinforcing its role as an alternative translation initiation site. In line, we have recently shown with rabbit reticulocyte lysates and HEK293T extracts that translation of Met11-Tau occurs from *in vitro*-synthesized tau mRNA transcripts (*43*). This mechanism is particularly intriguing, as dysregulated translation initiation is increasingly recognized in human diseases (*44, 45*). For instance, the recognition of alternative translation initiation sites (ATIS) by the ribosome producing N-terminally truncated variants have been linked to pathological outcomes (*46–48*), such as insulin resistance for using an ATIS at the Met-14 in Caveolin-2β (*49*). Our discovery of AcMet11-Tau suggests that non-canonical translational regulation may contribute to AD pathogenesis, warranting further investigation into the molecular mechanisms governing this process.

Moreover, alteration of the N-terminal acetylation pattern is also reported to be involved in human diseases (*21, 22, 50*). One key critical role of this prevalent protein modification is the regulation of protein stability; acetylation can protect proteins from degradation, thereby influencing protein half-life (*51*). Additionally, this modification has been shown to affect protein conformation and folding (*21*). For instance, in proteins associated with neurodegenerative diseases, such as α-synuclein, N-terminal acetylation influences their propensity to misfold and aggregate (*50*). For AcMet11-Tau, the functional consequences of N-α-acetylation remain to be fully elucidated. It is plausible that this modification enhances protein stability, protecting AcMet11-Tau from degradation and prolonging its half-life, or alters its folding, thereby promoting aggregation and seeding of full-length Tau as previously described (52).

The early detection of AcMet11-Tau in both transgenic mice and AD patient brains suggests an initiating role in disease progression. Our *in vivo* functional experiments further demonstrate that expression of AcMet11-Tau potentiates Tau pathology, while its targeted neutralization mitigates Tau pathology and cognitive decline. These results are consistent with prior reports, based on either expression or passive immunotherapy, showing that N-terminally truncated Tau variants contribute to Tau pathology development *in vivo* (*8, 53, 54*).

From a translational perspective, AcMet11-Tau holds dual potential as biomarker and therapeutic target. Tau immunotherapy is a promising avenue for AD treatment (*55–57*). Even the mechanisms underlying the beneficial effect of Tau-based immunotherapy are still to be determined, passive immunotherapies in rodent models of tauopathies have shown that monoclonal antibodies injected via the systemic pathway can cross the blood-brain barrier and bind to Tau-targeted proteins (*58, 59*). However, a major challenge lies in selectively targeting pathological Tau species without disrupting normal Tau function. Our findings address this challenge: AcMet11-Tau is exclusively detected in pathological contexts, and our specific antibody, 2H2D11, effectively binds this new variant without cross-reacting with physiological Tau. This selectivity, combined with the antibody’s specificity demonstrated efficacy in reducing pathology and cognitive deficits, underscores the therapeutic promise of AcMet11-Tau targeting.

Together, our findings identify a previously overlooked AD-relevant Tau species with dual potential as a biomarker and a novel therapeutic target. Further studies are required to assess the utility of AcMet11-Tau as a fluid or imaging biomarker. Ultrasensitive assays are needed to evaluate AcMet11-Tau as a fluid or imaging biomarker, in line with revised AD diagnostic criteria (*60*). The selective association of AcMet11-Tau with AD needs to be demonstrated by analysing more primary tauopathies. In parallel, additional research is needed to elucidate the molecular mechanisms underlying the production and pathogenicity of AcMet11-Tau, as it could also represent a therapeutic target. Finally, studies should assess the safety and efficacy of AcMet11-Tau-targeted immunotherapies in human trials.

## MATERIALS AND METHODS

### Study design

The primary aim of this study was to characterize AcMet11-Tau, a previously overlooked truncated Tau protein that we identified as an N-α-acetylated form in human brain by proteomic approach. To do this, it was necessary to get first a specific monoclonal antibody with no cross reactivity with full-length Tau proteins. This immunological tool was then used to examine brain tissues samples from transgenic mouse model of Tau pathology and human brains. Data analysis indicated that AcMet11-Tau may be implicated in Tau pathology development. To address this hypothesis, we subsequently performed functional studies in the transgenic mice.”

Two lines of Thy-Tau transgenic mice (described in mice section, bellow) were used in this study. Thy-Tau22 line is a well-established model for investigating modifiers of Tau pathology and their impact on memory. These mice develop progressive hippocampal Tau pathology before 3 months of age, with memory deficits developing from 6-7 months. In this model, Tau pathology has been reported to lack sexual dimorphism. This transgenic line was then used for IP passive immunizations.

Thy-Tau30 line exhibits a more rapid disease course, with widespread neurofibrillary tangle accumulation, axonopathy and severe motor deficits, which can confound memory assessments. Accordingly, this line was used only in experiments that did not involve behavioral testing.

Mice were IP immunized in statistically valid number of mice (n = 10–15 per group), based on prior experience with Thy-Tau22 model. Both male and female mice were included in the immunization groups. Behavioral assessments were performed in males only, while Tau pathology analyses were performed on pooled data from both males and females.

The selection of human brain areas in Fig.4 was based on both tissue availability and their relevance to the known spatiotemporal progression of tau pathology. Brain extracts were analyzed by Western blot with antibodies against Tau-Cter and Tau-pSer396 (fig. S2). One patient (C23, table. S1), whose biochemical profile matched the Braak stage III–IV group, was reassigned to this group for Elisa analyses despite a neuropathological diagnosis of NFT at Braak stage II. Additional points related to the study design are presented in other sections of the manuscript or in the supplementary materials.

### Human brain tissue

Frozen brain tissue samples from patients diagnosed with various tauopathies-Alzheimer’s disease (AD), Pick’s disease (PiD), and progressive supranuclear palsy (PSP)-as well as from neurologically healthy individuals, were obtained from the Lille NeuroBank collection (Centre de Ressources Biologiques du CHU de Lille). The Lille NeuroBank has been declared to the French Research Ministry by the Lille University Hospital (CHU-Lille) on August 14, 2008 under the reference DC-2000-642 and fulfills the criteria defined by French Law regarding biological resources, including informed consent; ethics review committee approval and data protection. The ethical review committee of Lille NeuroBank approved the study. The samples used in this study are listed in table S1.

### Mice

For primary neuronal culture, gestating C57Bl6/J mice were purchased from Janvier Laboratories (Le Genest-Saint-Isle, France). Thy-Tau transgenic mice lines (Thy-Tau22 and Thy-Tau30), C57Bl6/J background, that develop with age neurofibrillary degeneration and memory deficits were generated by the overexpression of a human Tau-412 isoform (1N4R) bearing two pro-aggregative mutations (G272V & P301S) under the control of the Thy1.2 neuronal promoter (*23*). Additional description is previously provided for Thy-Tau 22 model (*24, 31*) and for Thy-Tau-30 (*26*). Non-transgenic wild-type littermates (WT) were used as controls. The animals were housed in a pathogen-free facility as group of 5 in ventilated cages, with *ad libitum* access to food and water and maintained on a 12h light/12h dark cycle and controlled temperature (20-22°C). The animals were maintained in compliance with European standards for the care and use of laboratory animals and all experimental protocols were approved by the CEEA75 ethical committee (12787-2015101320441671v9).

### Tau plasmid and lentiviral vectors

Plasmid vectors carrying cDNA for Full-length Tau (Tau-412 isoform, Uniprot P10636-7) and Met11-Tau (in the same isoform context than FL-Tau) were generated using the In-Fusion cloning Kit (Clontech), and PCR primers were designed to clone inserts into the EcoRI site of pcDNA4/TO (Invitrogen). Each cDNA fragment was amplified by PCR (DyNAzymeTM EXT DNA polymerase, New England BioLabs) from pcDNA3.1-Tau4R (*61*). Forward primers were designed to contain the Kozak consensus sequence and are as follows: For Tau-412, 5’-CAGTGTGGTGGAATTCGCCACCATGGCTGAGCCCCGCCAGGAGTT-3’; For Met11-Tau, 5’-CAGTGTGGTGGAATTCGCCACCATGGAAGATCACGCTGGGACGT-3’. The reverse primer for both amplifications was: 5’-GATATCTGCAGAATTCTCACAAACCCTGCTTGGCCAGGGAGGCA-3’. Lentiviral vectors (Lv) with neuronal tropism (*62*), carrying cDNA for Full-length Tau and Met11-Tau were generated, produced and their titer determined as previously detailed (*25*); the titer was determined using the HIV-1 capsid protein p24 (Gentaur BVBA, Paris, France). DNA sequencing was carried out for constructs validation.

### Tau cell lines

SH-SYSY cells that constitutively express tetracycline repressor (*63*) were transfected with Tau plasmid constructs using ExGen500 (Euromedex, France) according to manufacturer’s instructions. Individual stable clones were generated following Zeocin selection (100 μg/ml), and those that exhibited the weakest basal expression of Tau were selected. The cell lines were maintained in Dulbecco’s modified Eagle’s medium (DMEM, Gibco BRL), supplemented with 10% fetal calf serum with pyruvate, 2mM L-glutamine, 50 units/ml penicillin/streptomycin and 1mM non-essential AA, in a 5% CO2 humidified incubator at 37°C. Tau expression is induced by Tetracycline at 1 μg/ml (Invitrogen).

## LC-MS/MS

AcMet11-Tau was identified using the experimental approaches previously detailed (11). Briefly, Tau species were enriched by immunoprecipitation from human occipital cortex brain (table S1), labeled with NHS-SS-Biotin to enrich the N-Terminal peptides, digested and analyzed by capillary liquid chromatography-tandem mass spectrometry (LC-MS/MS). The raw data were processed as described (*11*) and the mass spectrometry proteomic data have been deposited to the ProteomeXchange Consortium (http://proteomecentral.proteomexchange.org) via the PRIDE partner repository (*64*) with the dataset identifier PXD001353 and DOI 10.6019/PXD001353.

### Generation of monoclonal antibodies

2H2D11 antibody was generated by immunization with AcMet11-Tau peptide corresponding to the human Tau sequence from Methionine at position 11 to Glycine at position 21, encoded by exon1 and shared by all human Tau isoforms. A cysteine residue has been added to the peptide C-terminus for the coupling to a Keyhole limpet hemocyanin (KLH). Immunizations and generation of hybridomas were carried out by Agro-Bio (La Ferté Saint-Aubin, France), in accordance with applicable regulations. The peptides were synthesized by standard chemical peptide synthesis and purity was analyzed by HPLC (Agro-Bio, France). Balb/c mice were immunized subcutaneously with 3 boosts at days 14, 45 and 63. The lymphocytes from the spleen of the mouse displaying the highest titer were then fused with NS1 myeloma cells, according to the method described in (*65*). Supernatants from the resulting hybridomas were screened in indirect Elisa against the following peptides: AcMet11-Tau, Met11-Tau and AcMet1-Tau (Fig. 1C); the latter peptide starts at Methionine 1 of human Tau with an N-α-terminal acetylation. Elisa screenings allowed selection of clones that specifically detect the AcMet11-Tau with a slight or without any cross-reactivity with either Met11-Tau or AcMet1-Tau peptides. Isotype and the type of light chain have been determined for the selected hybridomas (The SBA Clonotyping™ System/HRP kit, # MBS678003). The hybridoma 2H2 was further subcloned and the clone that produces 2H2D11 antibody (IgG2a, Kappa) was selected. The specificity of 2H2D11 towards N-α-terminally acetylated methionine11 of Tau protein was consistently validated by Elisa, western blotting and immunohistochemistry.

An antibody against total Tau proteins was also generated by the same procedure, using Met11-Tau peptide (MEDHAGTYGLG) coupled to KLH for immunization. We selected the clone 7C12/E12 (IgG1, Kappa) that recognizes all the 3 Tau-peptides.

### Hybridoma cell culture for antibodies production

Hybridoma cell culture was performed in the 2-compartment bioreactor flasks CELLine CL 1000 (90-005, Integra Biosciences, France), using hybridoma serum free medium (SFM, 12045076, Gibco BRL) supplemented with 2mM L-glutamine and 50 units/ml penicillin/streptomycin, and maintained in a 5% CO2 humidified incubator at 37°C. High cell density culture was performed over a period of 20-30 days as previously described (*66*) with minor modifications. Hybridoma cells (2.5 x 10^7^ in 15 ml) were seeded in the cell compartment and the nutrient compartment was filled with 500 ml of the cell culture medium. The cell compartment medium harvest/replenish and nutrient medium exchange were performed every 3 days. The harvest from cell compartment was centrifuged at 250 x g for 5 min and the supernatant was then filtered with a 0.22 μM filter and kept frozen at −20°C until use for antibodies purification. Cell viability was checked during each cell compartment medium harvest by Trypan blue dye exclusion, using a cell counting chamber (87144F, Kova International, USA).

Monoclonal antibodies were purified from hybridoma culture supernatants (pool of 5-8 cell compartment medium harvests; see above) using an AKTA® FPLC system; the protocol is detailed in supplementary materials.

### Elisa assays

The detailed protocols for Indirect and sandwich Elisa are provided in supplementary materials.

### Stereotaxic injections

One-month-old heterozygous Thy-Tau30 transgenic mice were anesthetized with intraperitoneal injection of Ketamine (100 mg/ kg) and Xylazine (20 mg/ kg) mix. The animals were positioned on a stereotactic device (David Kopf Instrument) and bilateral injections were performed into CA1 region of the hippocampus, at the following coordinates with respect to the bregma: antero-posterior −2,5mm, medial-lateral −1mm (right side) and +1 (left side) and dorso-ventral −1.8mm. Equivalent amounts of Lentiviral vectors (445 ng of p24) or PBS (2.5 µl) were injected at a rate of 0.25 µl/min, using a 10 µl glass syringe with a fixed needle (Hamilton; Dutscher, Brumath, France). Two months post-injections, mice were deeply anesthetized with pentobarbital sodium (50 mg/kg, i.p.), then transcardially perfused, first with cold NaCl (0.9%) and then with 4% paraformaldehyde in 0.1 mol/L phosphate-buffered saline (pH 7.4) during 20 min. Brains were post-fixed during 1 day in 4% paraformaldehyde, then incubated in 20% sucrose for 24 hours, frozen 1min in Isopentane at −40°C then kept at −80°C until use.

The same procedure was used for intracerebral injection of monoclonal antibodies. Thy-Tau30 mice (3.5-month-old) received bilateral stereotaxic injections with 2 µg (1 µl) of 2H2D11 or control IgG (see below) into the hippocampus (coordinates relative to Bregma: AP: −2.5, ML: ±1, DV: −2.3). Five days post-injection, brains were processed as described above.

### Passive immunotherapy (fig. S4)

Repeated intraperitoneal injections (IP) of 2H2D11 antibody (IgG2a isotype) or control IgG (IgG2a isotype control, purified from B69 hybridoma (ATCC^®^ HB-9437^™^)) were performed in heterozygous Thy-Tau22 at the dose of 10 mg/kg. In the same experimental procedure, littermate WT mice were injected with either PBS or control IgG. Mice have received their first IP at 3 months of age and then every 2 weeks until behavioral evaluation at 7 months of age. Mice received a last injection one week before sacrifice (at 8 months of age). The animals were sacrificed by cervical dislocation and brains removed. The right hemispheres were post-fixed for 7 days in 4% paraformaldehyde, then incubated in 20% sucrose for 24 hours and kept frozen at −80°C until use. The left hemispheres were used to dissect out hippocampus by using a coronal acrylic slicer (Delta Microscopies) at 4°C and stored at −80°C for biochemical analyses.

### Immunohistochemistry (IHC)

Mice serial free-floating brain coronal sections (40µm) were obtained using a cryostat (Leica Microsystems GmbH, Germany) and kept in a PBS-azide (0.2 %) at 4°C until use. Sections of interest were washed with PBS-Triton (0.2%), treated for 30 min with 0.3% H2O2 and nonspecific binding was blocked with MOM (Mouse IgG blocking reagent) or horse serum (1/100 in PBS; Vector Laboratories) for 1 hour. The sections were then incubated with the primary antibody (table S2) in PBS-Triton 0.2% overnight at 4°C. After 3 x 10 min washes, labeling was amplified using a biotinylated anti-mouse or anti-rabbit IgG (1/400 in PBS-Triton 0.2%; Vector Laboratories) for 1 hour, followed by the ABC kit (1/400 in PBS; Vector Laboratories), and labeling was completed using 0.5 mg/ml DAB (Vector Laboratories) in 50 mmol/l Tris-HCl, pH 7.6, containing 0.075% H2O2. Brain sections were mounted on SuperFrost slides, dehydrated through a graded series of alcohol and toluene, and then mounted with Vectamount medium (#H5000; Vector Laboratories) and coverslips (#LCO2460M; Labelians) for microscopic analysis. Tissue sections were digitized using an automated slide scanner (Zeiss AXIOSCAN Z1) coupled with a high-resolution microscope (x 20 objective). The resulting virtual images were initially saved in CZI format and then converted to TIFF format using ZEN imaging software for export and further analysis. Quantification of immunostaining was performed using ImageJ software. Depending on the analysis type, either the number of immunopositive cells or the percentage of labeled surface area was quantified. Briefly, images were converted to 8-bit grayscale, and a thresholding procedure was applied to isolate the signal corresponding to antibody staining. This allowed for the measurement of either the number of positive events (cells) or the percentage of the area stained. For each section, the total surface area of the hippocampus was also measured and recorded.

### Immunofluorescence

Mice brain sections were washed with PBS-Triton 0.2%, and nonspecific binding was blocked through incubation with MOM (1/100 in PBS-Triton 0,2%; Vector Laboratories) for 1 hour. For co-labeling, the sections were incubated with the first primary antibody in PBS-Triton 0.2% overnight at 4°C. After 3 washes (10 min), the sections were incubated with the second primary antibody in PBS-Triton 0.2% overnight at 4°C. Sections were then washed and incubated with the two secondary antibodies (Invitrogen) coupled to either Alexa 468 or Alexa 588 (1/1000 in PBS) for 1 hour at room temperature. After 3 x 10 min washes, the sections were mounted with Vectashield containing DAPI to label the nuclei (Vector Laboratories). Confocal microscopy was performed on a Zeiss LSM 710 inverted confocal microscope (x63 oil objective). Images were collected in the z direction at 1μm or 0.8 μm intervals.

### Peptide blocking experiments

Before proceeding to Western Blot or IHC, 2H2/D11 antibody was incubated with agitation overnight at 4°C, in blocking buffer without or with excess of the blocking peptide (molar ratio: 1/50). Thereafter, the antibody samples were used to perform staining protocol as described in Western Blotting and IHC sections.

### Protein extractions

Detailed procedures for preparation of total proteins from cells and brain samples are provided in supplementary materials.

### Western Blotting

For total proteins, extracts were standardized at 1mg/ml with LDS 2X supplemented with a reducing agent (Invitrogen) and denatured at 100°C for 10 min. Proteins were then separated with SDS-PAGE using precast 4-12% Bis-Tris NuPage Novex gels (Invitrogen). Proteins were transferred to 0.45 µM nitrocellulose membranes (AmershamTM Hybond ECL), which were saturated with 5% non-fat milk or bovine serum albumin (Sigma) in TNT buffer (140 mM NaCl, 0.5% Tween20, 15mM Tris, pH 7.4), according to the primary antibody. Membranes were then incubated with the primary antibody overnight at 4°C, washed with TNT buffer 3 times for 10 min, incubated with the HRP-conjugated secondary antibody (Vector) for 45 min at RT and washed again. Immunolabeling was visualized using chemiluminescence kit <u>(ECL #RPN2106;</u> Amersham) and LAS-4000 acquisition system (Fujifilm). The antibodies list is given in table S2.

### Behavioral Tests

To avoid bias, animals were randomly allocated to experimental groups, and behavioral testing was performed by investigators blinded to treatment conditions. The detailed procedures for Actimetry, Elevated Plus Maze, Y-Maze and Barnes Maze are provided in the supplementary materials.

### RNA Extraction and Quantitative RT-PCR

Total RNA was extracted from dissected mouse hippocampi using the RNeasy Lipid Tissue Mini Kit (Qiagen), according to the manufacturer’s instructions. RNA concentrations were determined using a NanoDrop ND-1000 spectrophotometer at 260 nm. RNA purity was assessed using A260/A280 and A260/A230 absorbance ratios. Extracted RNA samples were stored at –80°C until use. Detailed protocol for RT-PCR is provided in supplementary materials.

### Statistical analysis

Statistical analyses were performed using Prism 10 software (GraphPad Software Inc., USA). Data are presented as mean ± standard error of the mean (SEM). The normality of data distribution within groups was assessed using the Shapiro–Wilk test. Depending on the number of experimental groups, comparisons were performed using either unpaired t-test or one-way ANOVA, followed by appropriate multiple comparison post hoc tests (Tukey’s HSD or Fisher’s LSD). Correlation analyses between continuous variables were carried out using Spearman’s rank correlation. For all statistical tests, a p-value < 0.05 was considered statistically significant. The specific details are indicated in the figure legends.

## Acknowledgments

We thank the staff of the different facilities of ‘Plateformes Lilloises en Biologie et Santé (PLBS)’–UAR 2014–US 41 (SOFP facility, *In vivo* and functional exploration platform and BiCeL), with special thanks to M. Tardivel and A. Bongiovanni for their assistance with microscopy acquisitions and analyses. We thank Sébastien Carrier, Espérance Pastouret and Anna Bogdanova for their help in animal genotyping. We also think Pr. P Davies for his generous gift of MC1 antibody. We are thankful to the patients and their families for their generous approval to use postmortem brain material for research. We also thank the neurologists from the CMRR (Centre Mémoire de Ressources et de Recherche, CHU Lille) and the neuropathologists from the CRB (Départements universitaires d’Anatomie Pathologique et d’Histologie, Centre de Ressources Biologiques, CHU Lille)

## Author contributions

Conceptualization: LB and MH, with the intellectual contribution from MD, FM and DB Methodology: SG, CLe, MD, SE, SL, FV, RC, SB, MH

Formal analysis: SG, CLe, MD, SE, SL, CLa, GC, YV, DB, MH Investigation: SG, CLe, MD, SE, SL, FV, KC, MH Visualization: SG, CLe, MD, SE, SL, CR, YV, DB, MH Resources: DB, VD, VBS, SS, LB

Funding acquisition: FM, LB, MH Project administration: MH Supervision: LB, MH

Writing – original draft: MD, LB, MH

Writing – review & editing: MD, RC, CLa, PJdC, KC, GC, SS, DB, FM, LB, MH

## Data and materials availability

All data and methods used in this study are available in the main text or the supplementary materials. The mass spectrometry proteomics data have been deposited to the ProteomeXchange Consortium (http://proteomecentral.proteomexchange.org) via the PRIDE partner repository (64) with the dataset identifier PXD001353 and DOI 10.6019/PXD001353. The VH and VL sequences of 2H2D11 antibody are available on request. Cell and animal models, as well as the antibodies generated in this study are available by contacting LB under MTA with Inserm.

