## Supplementary material for "N-terminally acetylated Met11-Tau: a new pathological truncated Tau species with functional relevance in Alzheimer disease": Materials and Methods Fig. S1 to Fig S6 Tables S1 to S3

### Supplementary Materials and Methods

#### Primary neuronal culture and infection with Lentiviral vectors

Primary cortical neurons were prepared from 15-days-old C57Bl6/J mouse embryos as previously described (67). Once dissociated and counted, cells were plated in poly-D-lysine (0.5 mg/ml, Sigma-Aldrich) and laminin (10 µg/ml, Sigma-Aldrich)-coated 6-well plates (800 000 cells per well) and maintained in a 5% CO2 humidified incubator at 37°C.

Lentiviral vector-based infections were performed at DIV11, as previously described (67); 400 ng of Lvs were added per well and 3 days later, the cells were washed once with phosphate-buffered saline and recovered in lysis buffer for WB analysis.

#### Purification of Monoclonal Antibodies using the AKTA® System

Monoclonal antibodies were purified from hybridoma culture supernatants (pool of 5-8 cell compartment medium harvests; see "Hybridoma cell culture for antibodies production" in Mat & Meth section) using an AKTA® FPLC system. This automated chromatography platform was equipped with a 5 mL HiTrap Protein A or Protein G affinity column (Cytiva). The column was first equilibrated with binding buffer (20 mM sodium phosphate, pH 7.0). Culture supernatant samples were injected onto the column, and after wash with binding buffer, elution is performed using 100 mM glycine-HCl, pH 2.7. Successive elution fractions (approximately 10–15 mL total volume per purification run) were collected and were analyzed for antibody presence by SDS-PAGE electrophoresis (NuPAGE Novex Bis-Tris 4–12% gels, Invitrogen). A 6 µL aliquot of each fraction was loaded onto the gel. Following migration, gels were stained with Coomassie Brilliant Blue G-250 (0.1% Coomassie in 50% ethanol and 10% acetic acid) and subsequently de-stained in a solution containing 7% acetic acid and 10% ethanol. This analysis allowed identification of elution fractions containing the purified antibody. Positive fractions were pooled and dialyzed against PBS using a 14 kDa dialysis tubing cellulose membrane (D9777-100FT, Sigma Aldrich), overnight at 4°C under agitation. The antibody solution was then quantified using the BCA assay kit (Pierce). If the final antibody concentration was below 1 mg/mL, a concentration step was performed using Amicon Ultra-15 centrifugal filter units with a 30 kDa molecular weight cut-off (Merck-Millipore). Two quality control steps were performed: 1) Integrity assessment by SDS-PAGE analysis of antibody samples treated or untreated with the reducing agent DTT (dithiothreitol), to confirm that the purification procedure did not induce antibody denaturation, 2) Functionality verification by Western blot, using protein lysates from cells expressing or not the target antigen. This confirmed the antigen-binding functionality of the purified antibody. If both quality controls were satisfactory, the antibody was considered suitable for downstream applications. For immunotherapy-related application, endotoxin levels were measured using the Pierce™ chromogenic endotoxin Quant Kit (Thermo Fisher Scientific). Endotoxin concentrations were found to be 0.66 EU/mL for the purified 2H2D11 antibody and 3.14 EU/mL for the control IgG. An endotoxin level of less than 10 EU/mL is recommended for preclinical studies involving *in vivo* antibody administration (*68*).

**Elisa**

Nunc 96-well microtiter plates (Maxisorp F8; Nunc, Inc.) were coated overnight at 4°C with 100ng/well of either AcMet11-Tau peptide, Met11-Tau peptide or AcMet1-Tau peptide in 50 mM NaHCO3, pH 9.6. After 3 washes with PBS containing 0.05% Tween (PBS-T), plates were blocked with 0.1% casein solution (PBS) at 37°C for 1 h, followed by incubation with hybridoma supernatants or purified antibodies; 2H2D11 or 7C12 (dilution in PBS containing 0.2% BSA) for 2 h at room temperature. After 3 washes with PBS-T, immunodetection was performed by using a goat anti-mouse IgG horseradish peroxidase-conjugated antibody (A3673; Sigma) at 1:4000 dilution in PBS-BSA 0.2%, at 37°C for 1 h. After 5 washes with PBS-T, detection was performed using Tetramethyl benzidine substrate (T3405, Sigma) for 30 min at room temperature; the assay was stopped with H2SO4 and absorbance was read at 450nm with (Multiskan Ascent spectrophotometer, Thermo Labsystems). All the peptides were synthesized by standard chemical peptide synthesis and purity was analyzed by HPLC (ProteoGenix, France).

**Sandwich Elisa**

Nunc 96-well plates (VWR) were coated with 100 µl (100 ng) of 2H2D11 antibody (for detection of AcMet11-Tau) or 7C12 antibody (detection of total Tau) in Carbonate buffer (NaHCO3 0.1M, Na2CO3 0.1M, pH 9.6) overnight at 4°C. The plates were subsequently blocked with a WASH1X buffer (INNOTEST hTau Ag kit, FUJIREBIO) containing 0.1% casein at 37°C for 1 hour and washed with WASH1X buffer 3 times. Protein samples were standardized at 1mg/ml and diluted in SAMPL DIL buffer (INNOTEST hTau Ag kit, FUJIREBIO). Protein samples and biotinylated

antibodies (HT7/BT2, INNOTEST hTau Ag kit, FUJIREBIO) were added and the plates were incubated at room temperature overnight. The wells were washed four times then incubated with Peroxidase-labeled streptavidin at room temperature for 30 min and washed four times. Detection was performed using Tetramethyl benzidine substrate for 30 min at room temperature; the assay was stopped with H2SO4 and absorbance was read with spetrophotometer (Multiskan Ascent, Thermo Labsystems) at 450nm.
For detection of Tau phosphorylated at Ser396, ELISA experiments were performed, accordingly to manufacturer's instructions, by using Human Tau [pS396] ELISA Kit (Invitrogen).

**Protein extractions**

Cells were washed with PBS and collected in ice-cold RIPA buffer (150 mM NaCl, 1% NP40, 0.5% sodium deoxycholate, 50 mM Tris-HCl, pH 8.0) completed with protease inhibitors cocktail (Complet mini EDTA-Free; Roche). After sonication (20 pulses at 40 Hz) and homogenization 30 min at 4°C under agitation, the supernatants were recovered after centrifugation at 12 000 × g for 10 minutes at 4˚C.
Brain tissue samples were processed following a standardized protocol. Approximately 100 mg of human brain tissue were homogenized by sonication (30 pulses at 40 Hz) in 1ml of cryopreservation buffer composed of 10 mM Tris-HCl and 0.32 M sucrose (pH 7.4), supplemented with a protease inhibitors cocktail (Complet mini EDTA-Free, Roche). For mice, 1/2 hippocampus was homogenized in 200 µl of cryopreservation buffer. Homogenates were stored at –80°C until use. For total protein extraction, an aliquot of the tissue suspension was mixed (1:1, v/v) with 2X RIPA ice-cold buffer, and protein extraction was carried out as described above. Protein concentrations were determined using the BCA Assay Kit (Pierce) and the extracts were kept at -80°C until use in Western blotting or ELISA

**Behavioral Tests**

***Actimetry:*** spontaneous locomotor activity was assessed using actimetry arenas (Bioseb, LE8816; 45 × 45 × 35 cm) lightened at 40 lux. Each arena was equipped with two infrared beam frames: one positioned 2 cm above the floor to detect horizontal movements, and another at 6 cm to assess vertical activity (rearing). Mice were habituated to the testing room for at least 30 minutes prior to recording. During the test session, each mouse was allowed to freely explore the arena for 10 minutes. Locomotor parameters including velocity, and total distance moved, were recorded using the Actitrack software (Bioseb, Vitrolles, France).

***Elevated Plus Maze (EPM):*** anxiety-like behavior was evaluated using the EPM, consisting of four arms (35 cm long × 5 cm wide), arranged in a plus configuration and elevated 60 cm above the floor. Two arms were open (no walls), and two were enclosed by 15 cm-high opaque walls. Mice were placed in the center of the maze and allowed to explore freely for 5 minutes. Locomotor activity and the time spent in each arm were recorded and analyzed using the EthoVision XT tracking system (Noldus, Wageningen, the Netherlands).

***Y-Maze:*** short-term spatial memory was assessed using the Y-maze, composed of three opaque arms (22 cm long × 6.4 cm wide × 15 cm high) arranged at 120° angles. Spatial cues were positioned on the surrounding walls to support spatial orientation. For each animal, the roles of the "start", "novel", and "other" arms were randomly assigned. In the learning phase, the novel arm was blocked by an opaque door. The mouse was placed in the start arm and allowed to explore the two accessible arms (start and other) for 5 minutes. After a 2-minute inter-trial interval in the home cage, the novel arm was opened, and the mouse was placed again in the start arm for the test phase, where it could explore all three arms for an additional 5 minutes (from the time the mouse first left the start arm). Time spent in each arm during both phases was recorded using EthoVision XT (Noldus, Wageningen, the Netherlands).

***Barnes Maze:*** spatial learning was assessed using the Barnes maze task. The maze consisted of a white circular PVC platform (120 cm diameter), placed on a swivel system in the center of the room, elevated 80 cm above the floor and brightly lit (370 lux). Forty equally spaced holes (5 cm in diameter) were arranged along the perimeter, 5 cm from the edge. Only one hole led to a dark escape box under the platform. Visual cues were placed on the walls surrounding the maze to facilitate spatial orientation. A camera positioned above the maze recorded animal behavior, which was analyzed using EthoVision XT (Noldus, Wageningen, the Netherlands). In the habituation phase (day 0), mice explored the maze for 2 minutes. If they failed to find the escape hole, they were gently guided toward it and allowed to remain inside for 30 seconds. The escape hole location was randomized for each mouse. In the acquisition phase (days 1-4), mice underwent four trials per day to learn the escape hole location. At the start of each trial, animals were placed under an opaque cylinder in the center of the platform for 10 seconds. Upon cylinder removal, mice had 3 minutes to locate the target hole. Animals that failed to do so on day 1 were guided to the hole and remained in the escape box for 30 seconds. Between trials, mice were returned to their home cage for a 15-minutes inter-trial interval. The platform was cleaned with 70% ethanol and rotated by 45° daily to prevent olfactory cue reliance. The order of testing was pseudo-randomized across animals to avoid experimental bias.

For the acquisition phase, the number of primary errors (number of times the mouse explores the other holes before finding the target one), total errors and distance moved were recorded.

**Quantitative RT-PCR**

Reverse transcription (RT) was performed using 1 µg of total RNA and the High-Capacity cDNA Reverse Transcription Kit (Applied Biosystems), in a T Gradient Biometra thermocycler, following the thermal conditions: 25°C for 10 s, 37°C for 2 h, and 85°C for 5 s. The resulting cDNAs were stored at –80°C until further analysis. Quantitative PCR (qPCR) was carried out on an Applied Biosystems Prism 7900 using a 1:20 dilution of cDNA and the Power SYBR™ Green PCR Master Mix (Applied Biosystems), or 1:10 dilution of cDNA and TaqMan gene expression master mix (Life technologies). The primers (table S3) were used at a final concentration of 0.1 nM. The qPCR program was as follows: 2 min at 50°C, 10 min at 95°C, followed by 40 cycles of 15 s at 95°C and 25 s at 60°C. A final step included 15 s at 95°C, followed by 1 min at 60°C, and a gradual temperature increase of 1°C increments up to 95°C to generate a melting curve and verify the specificity of the amplification product.

The peptidyl-prolyl isomerase A (PPIA), a ubiquitously expressed housekeeping gene with stable expression under experimental conditions, was used as the endogenous reference gene for normalization. All reactions were performed in triplicate, and relative gene expression was calculated using the comparative Ct method (ΔΔCt).

### Supplementary figures and tables

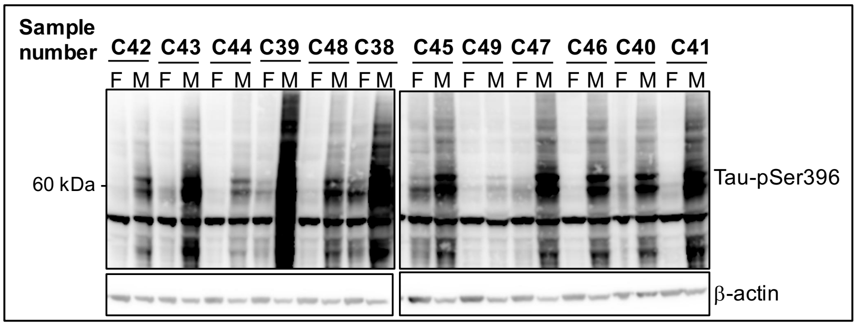


**Fig. S1**. **Biochemical characterization of Tau pathology in PSP Brain samples.** Protein extracts from frontal (F) and mesencephalon (M) brain regions were analyzed by western blot using an antibody against phosphorylated Tau proteins (Tau-pSer396); for each individual, typical pattern of Tau pathology is displayed mainly in mesencephalon than in frontal cortex. β-actin was used as loading control.

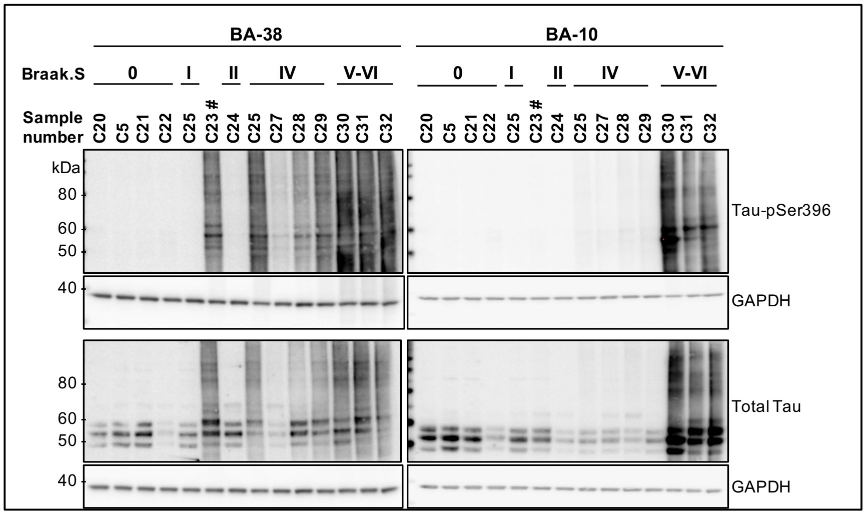


**Fig. S2. Biochemical characterization of Tau pathology regarding Braak stages.** Brain protein extracts from Borodmann areas 38 (BA-38) and 10 (BA-10) of individuals spanning Braak stages 0 to VI (neuropathological diagnosis), using Tau-pSer396 and Total Tau (C-ter) antibodies. # Individual whose biochemical profile matched the Braak stage III–IV group, was reassigned in this study to this group despite a neuropathological diagnosis of NFT at Braak stage II. GAPDH was used as a loading control.

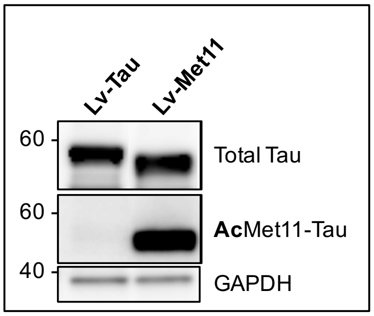


**Fig. S3**. **Validation of lentiviral vectors batches in primary neuronal cells.** Western blot analysis of protein extracts from neuronal cells, 3 days post-lentiviral vectors infections. GAPDH was used as loading control.

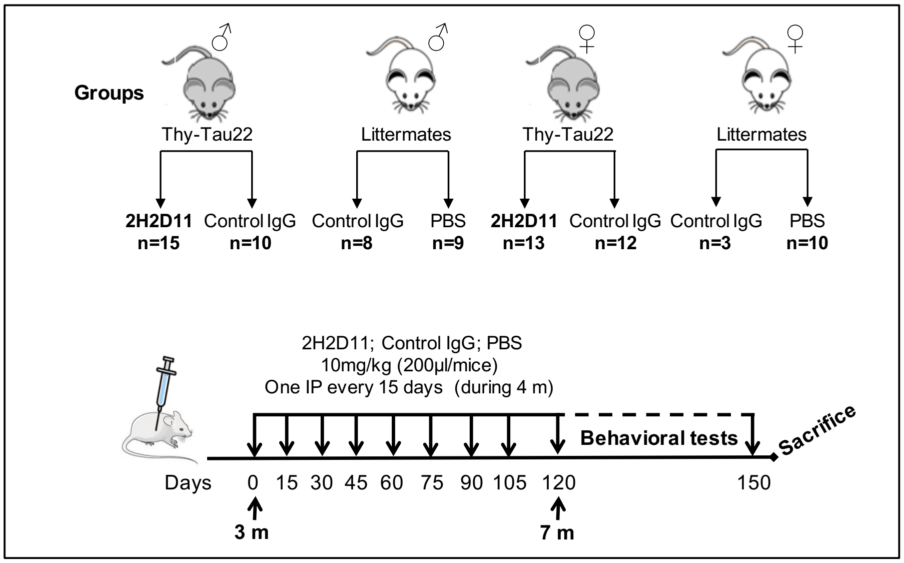


**Fig. S4. Experimental design of passive immunotherapy.** Experimental design of passive immunotherapy against AcMet11-Tau in Thy-Tau22 mice. Repeated intraperitoneal injections (IP) of the monoclonal antibody directed against AcMet11-Tau (2H2D11 antibody; IgG2a isotype) as well as with a control IgG antibody (IgG2a isotype, purified from B69 hybridoma (ATCC® HB-9437™)) were performed in heterozygous Thy-Tau22 males (10 mg antibody/kg). In the same experimental procedure, littermate WT mice groups were injected with either PBS (n=9) or control IgG (n=8), to get a baseline in behavioral evaluations (pooled as WT control group in behavioral tests). Experimental groups made of females were also included. Mice have received their first IP at 3 months of age and then every 2 weeks until behavioral evaluation at 7 months of age. Mice received a last injection one week before sacrifice (at 8 months of age). Body weights were measured monthly.

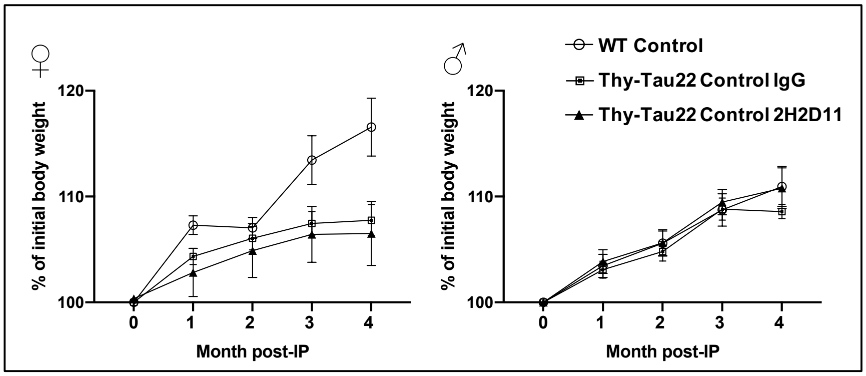


**Fig. S5**. **Mouse body weight during immunization.** Body monitoring curves showed no significant difference in body weight over the period of 4 months post intra-peritoneal injections (IP). Data are presented as % of initial body weight (mean ±SEM, n= 10-17/grp, two-away ANOVA followed by a post-hoc HSD Tukey test).

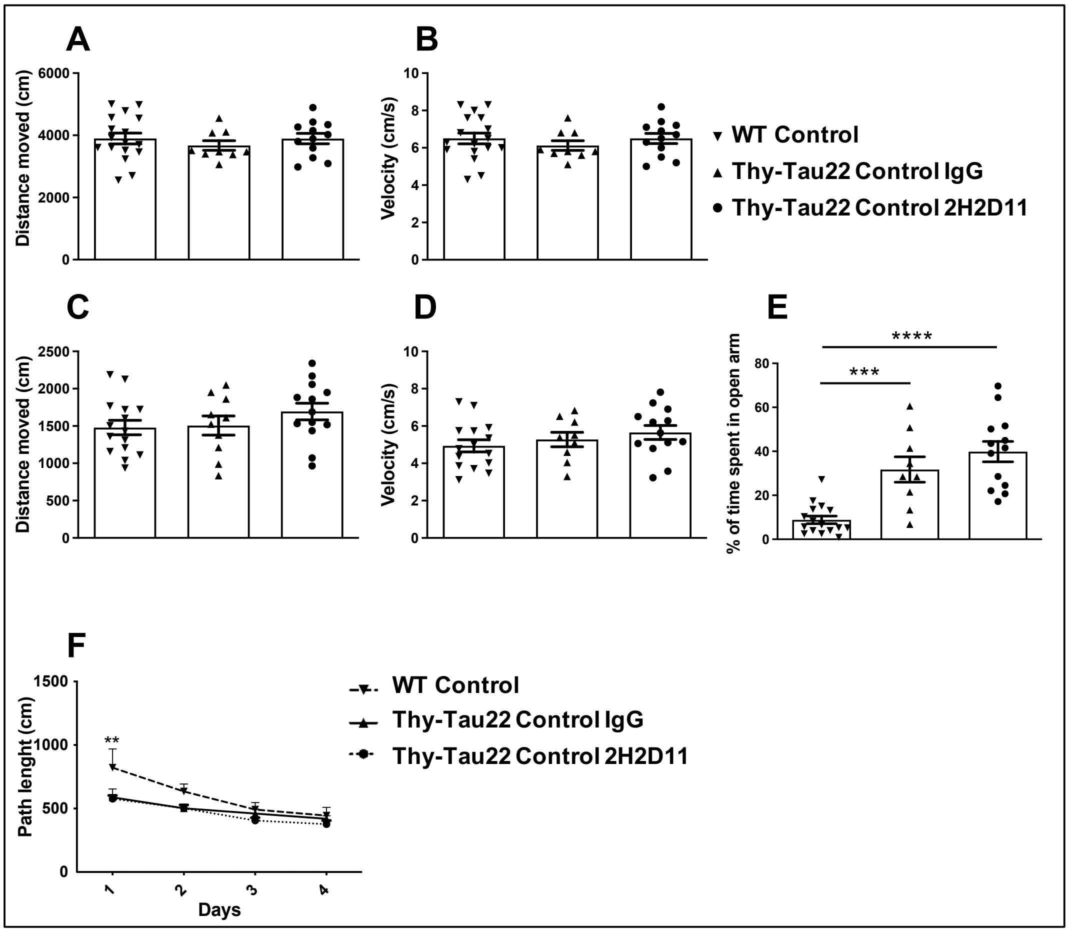


**Fig. S6. Locomotor activity and anxiety-like behavior.** (**A, B**) no difference was observed in Actimetry test regarding the distance moved and velocity. (**C to E)** no difference was observed in the EPM test regarding the distance moved (C) and velocity (D). The time spent in the open arms (E) is as expected significantly different between the WT mice and Thy-Tau22 mice treated with control IgG but no difference was observed between Thy-Tau22 mice treated with 2H2D11 antibody and control IgG. ****p<0.0001, ***p=0.003, one-way ANOVA followed by a post-hoc HSD Tukey test. (**F**) path length during the 4 days of acquisition in the Barnes maze. *p <0.05, two-way ANOVA followed by a post-hoc HSD Tukey test. Data are presented as mean ±SEM (n=10-17/group).

**Table S1. Summary of human brain tissues.**

| Sample number | Pathology | Gender | Age | Application |
|---|---|---|---|---|
| C1 | Control | M | 29 | Elisa (Fig. 3, A to D) |
| C2 | Control | M | 61 | Elisa (Fig. 3, A to H) |
| C3 | Control | M | 75 | Elisa (Fig. 3, A to D) |
| C4 | Control | F | 80 | Elisa (Fig. 3, A to D) |
| C5 | Control (Braak 0) | M | 22 | Elisa (Fig. 3, A to D)/ Fig. 4/fig. S2 |
| C6 | control | F | 82 | Elisa (Fig. 3, A to H) |
| **C7** | ***(Braak III)** | M | 76 | LC-MS/MS (Fig. 1A) |
| C8 | AD (Braak IV) | M | 84 | Elisa (Fig. 3, A to F) |
| C9 | AD (Braak VI) | M | 76 | Elisa (Fig. 3, C and D) |
| C10 | AD (Braak VI) | M | 86 | Elisa (Fig. 3, A to D) |
| C11 | AD (Braak V) | F | 76 | Elisa (Fig. 3, A to D) |
| C12 | AD (Braak V) | M | 77 | Elisa (Fig. 3, A to F) |
| C13 | AD (Braak V) | M | 62 | Elisa (Fig. 3, A to F) |
| C14 | AD (Braak VI) | F | 86 | Elisa (Fig. 3, E and F) |
| C15 | AD (Braak IV) | F | 91 | Elisa (Fig. 3, C and D) |
| C16 | AD (Braak VI) | M | 57 | Elisa (Fig. 3, A to D) |
| C17 | AD (Braak VI) | F | 73 | Elisa (Fig. 3, A to F) |
| C19 | AD (Braak VI) | F | 72 | Elisa (Fig. 3, A to F) |
| C20 | Control (Braak 0) | M | 58 | Elisa (Fig. 4)/fig. S2 |
| C21 | Control (Braak 0) | M | 41 | Elisa (Fig. 4)/fig. S2 |
| C22 | Control (Braak 0) | M | 78 | Elisa (Fig. 4)/fig. S2 |
| **C23** | ***(Braak III-IV)** | F | 89 | Elisa (Fig. 4)/fig. S2 |
| **C24** | *** (Braak II)** | F | 81 | Elisa (Fig. 4)/fig. S2 |
| **C25** | ***(Braak I)** | M | 59 | Elisa (Fig. 4)/fig. S2 |
| C26 | AD (Braak IV) | M | 84 | Elisa (Fig. 4)/fig. S2 |
| C27 | AD (Braak IV) | M | 77 | Elisa (Fig. 4)/fig. S2 |
| C28 | AD (Braak IV) | F | 88 | Elisa (Fig. 4)/fig. S2 |
| C29 | AD (Braak IV) | F | 86 | Elisa (Fig. 4)/fig. S2 |
| C30 | AD (Braak VI) | F | 81 | Elisa (Fig. 4)/fig. S2 |
| C31 | AD (Braak VI) | M | 65 | Elisa (Fig. 4)/fig. S2 |
| C32 | AD (Braak V) | F | 69 | Elisa (Fig. 4)/fig. S2 |
| C33 | Pick | F | 85 | Elisa (Fig. 3, E and F) |
| C34 | Pick | M | 57 | Elisa (Fig. 3, E and F) |
| C35 | Pick | M | 71 | Elisa (Fig. 3, E and F) |
| C36 | Pick | F | 68 | Elisa (Fig. 3, E and F) |
| C37 | Pick | F | 68 | Elisa (Fig. 3, E and F) |
| C38 | PSP | M | 57 | Elisa (Fig. 3, E to H)/fig. S1 |
| C39 | PSP | F | 77 | Elisa (Fig. 3, E to H)/fig. S1 |
| C40 | PSP | M | 75 | Elisa (Fig. 3, E to H)/fig. S1 |
| C41 | PSP | M | 82 | Elisa (Fig. 3, E to H)/fig. S1 |
| C42 | PSP | F | 82 | Elisa (Fig. 3, E to H)/fig. S1 |
| C43 | PSP | F | 77 | Elisa (Fig. 3, E to H)/fig. S1 |
| C44 | PSP | M | 85 | Elisa (Fig. 3, E to H)/fig. S1 |
| C45 | PSP | M | 87 | Elisa (Fig. 3, E to H)/fig. S1 |
| C46 | PSP | F | 79 | Elisa (Fig. 3, E to H)/fig. S1 |
| C47 | PSP | F | 74 | Elisa (Fig. 3, E to H)/fig. S1 |
| C48 | PSP | F | 77 | Elisa (Fig. 3, E to H)/fig. S1 |
| C49 | PSP | M | 90 | Elisa (Fig. 3, E to H)/fig. S1 |

For controls and AD cases, Braak staging was categorized by experienced neuropathologists based on histological analysis and the Braak classification system (69). Patients were grouped according to the stage of neurofibrillary degeneration (NFT pathology) as follows: Braak stage 0 (no detectable neurofibrillary lesions), Braak stages I–II (early-stage NFTs), Braak stages III–IV (moderate NFT pathology), Braak stages V–VI (severe NFT pathology). Braak stages I–III indicate early to moderate neurofibrillary pathology (involving the hippocampus, limbic regions, and subsequently temporal areas), yet this remains insufficient to support a definitive diagnosis of Alzheimer's disease (AD). *Importantly, cases with only Braak stages I–III can hardly be labeled as "Alzheimer's disease" since neurofibrillary pathology at these stages is not sufficient to establish a definitive diagnosis. These cases are therefore better described as exhibiting early neurofibrillary pathology or compatible with preclinical AD, rather than confirmed AD. Most individuals in the Braak 0 and I–II groups had no clinical history of cognitive impairment or dementia at the time of death. Some individuals in the Braak III–IV group exhibited mild cognitive symptoms. Patients classified as Braak stage V–VI were selected based on documented clinical histories consistent with typical Alzheimer's disease progression. After tissue processing, samples were analyzed by Western blot to confirm NFT pathology and staging (5).

### Table S2. Antibodies used in this study.

| Antibody | Specificity (Epitope) | Species | Applicaion (Dilution) | Source |
|---|---|---|---|---|
| Tau-Cter | Total-Tau (aa 426-441) | Rabbit, polyclonal | WB (1/10000) | Home-made |
| 7B1 | Total-Tau (aa 162-175) | Mouse, monoclonal | IHC (1/400; 2.5 μg/ml) | Home-made |
| MC1 | Pathological Tau conformati (aa 312-322) | Mouse, monoclonal | IHC (1/1000) | Generous gift from P. Davies (Gisha et al., 1999) |
| AT100 | PHF-Tau (pThr212/pSer214) | Mouse, monoclonal | WB (1/1000), IHC (1/500) | Invitrogen #MN1060 |
| Tau p-Ser199 | Tau (Ser199) | Rabbit, polyclonal | IHC (1/500) | 4BioDx #4BDX-1502 |
| Tau p-Ser396 | Tau (pSer396) | Rabbit, polyclonal | WB (1/10000) | Invitrogen #44-752G |
| Tau p-Ser422 | Tau (pSer422) | Rabbit, polyclonal | IHC (1/100) | Invitrogen #44-764G |
| 7C12 | Total-Tau (aa 11-20) | Mouse, monoclonal | WB and indirect ELISA (1/5000:0.2 μg/ml) | Generated in this study |
| 2H2D11 | Truncated AcMet11-Tau (α-acetylated Met11) | Mouse, monoclonal | WB and indirect ELISA (1/5000; 0.2 μg/ml), IHC (1/200; 5 μg/ml), Immunotherapy | Generated in this study |
| B 69 | Infectious Bursal Disease virus | Mouse, monoclonal | Immunotherapy | Purified in our Lab from: B69 ATCC HB-9437 hybridoma |
| GAPDH | Mouse GAPDH (FL1-335) | Rabbit, polyclonal | WB (1/10000) | Sigma-Aldrich #G9545 |
| NSE | the center region | Rabbit, polyclonal | WB (1/50000) | Gene Tex |
| Anti-β-Actin | N-terminus | Mouse, monoclonal | WB (1/1000) | Sigma-Aldrich #A5441 |

### Table S3. Primers used in qPCR.

| Designed primers (Syber Green) | | | | |
|---|---|---|---|---|
| **Name** | **Accession number** | **Forward primer** | **Reverse primer** | **Amplicon (bp)** |
| CD68 | NM_009853.1 | GACCTACATCAGAGCCCGAGT | CGCCATGAATGTCCACTG | 95 |
| Clec7A | NM_020008.2 | ATGGTTCTGGGAGGATGGAT | GCTTTCCTGGGGAGCTGTAT | 72 |
| Itgax | NM_0211334.2 | ATGGAGCCTCAAGACAGGAC | GGATCTGGGATGCTGAAATC | 62 |
| GFAP | NM_001131020.1 | CGCGAACAGGAAGAGCGCCA | GTGGCGGGCCATCTCCTCCT | 104 |
| TLR2 | NM_011905.3 | GGGGCTTCACTTCTCTGCTT | AGCATCCTCTGCGATTTGACG | 110 |
| Cyclophilin A (PPIA) | NM_008907.1 | AGCATACAGGTCCTGGCATC | TTCACCTTCCCAAAGACCAC | 126 |

| Taqman probes | | | | |
|---|---|---|---|---|
| Name | Accession number | Assay ID | Assay Design | Amplicon (bp) |
| C1qa | NM_007572.2 | Mm00432142_m1 | Probe spans exons | 80 |
| Cyclophilin A (PPIA) | NM_008907.1 | Mm02342430_g1 | Probe spans exons | 148 |
